# Understanding the origins of loss of protein function by analyzing the effects of thousands of variants on activity and abundance

**DOI:** 10.1101/2020.09.28.317040

**Authors:** Matteo Cagiada, Kristoffer E. Johansson, Audrone Valanciute, Sofie V. Nielsen, Rasmus Hartmann-Petersen, Jun J. Yang, Douglas M. Fowler, Amelie Stein, Kresten Lindorff-Larsen

**Author notes:** **For correspondence:** (KLL).

## Abstract

Understanding and predicting how amino acid substitutions affect proteins is key to our basic understanding of protein function and evolution. Amino acid changes may affect protein function in a number of ways including direct perturbations of activity or indirect effects on protein folding and stability. We have analysed 6749 experimentally determined variant effects from multiplexed assays on abundance and activity in two proteins (NUDT15 and PTEN) to quantify these effects, and find that a third of the variants cause loss of function, and about half of loss-of-function variants also have low cellular abundance. We analyse the structural and mechanistic origins of loss of function, and use the experimental data to find residues important for enzymatic activity. We performed computational analyses of protein stability and evolutionary conservation and show how we may predict positions where variants cause loss of activity or abundance. In this way, our results link thermodynamic stability and evolutionary conservation to experimental studies of different properties of protein fitness landscapes.

## Introduction

Mutational analysis of proteins have provided us with a wealth of information about the molecular interactions that stabilize proteins and govern their functions (***Fersht, 1999***). This information has in turn enabled us to engineer proteins with improved activities and stability (***Goldenzweig and Fleishman, 2018***), to better understand how mutations cause disease (***Stein et al., 2019***), and help elucidate the role of protein stability in evolution (***DePristo et al., 2005***; ***Echave et al., 2016***).

Computational analyses of missense variants in genetic diseases have suggested that loss of function via loss of protein stability is a major cause of disease (***Wang and Moult, 2001***; ***Ferrer-Costa et al., 2002***; ***Steward et al., 2003***; ***Yue et al., 2005***; ***Casadio et al., 2011***; ***Gao et al., 2015***; ***Stein et al., 2019***) because unstable proteins either aggregate or become targets for the cell’s protein quality control apparatus and are degraded (***Nielsen et al., 2020***). Indeed, cellular studies of disease-causing variants in a number of genes have shown that many variants are degraded in the cell (***Meacham et al., 2001***; ***Yaguchi et al., 2004***; ***Olzmann et al., 2004***; ***Ron and Horowitz, 2005***; ***Yang et al., 2011,2013***; ***Arlow et al., 2013***; ***Nielsen et al., 2017***; ***Chen et al., 2017***; ***Matreyek et al., 2018***; ***Scheller et al., 2019***; ***Abildgaard et al., 2019***; ***Suiter et al., 2020***). For this reason, several methods for predicting and understanding disease-causing variants include predictions of changes in protein stability (***Yue et al., 2005***; ***De Baets et al., 2012***; ***Casadio et al., 2011***; ***Ancien et al., 2018***; ***Wagih et al., 2018***; ***Gerasimavicius et al., 2020***).

While stability-based predictions can be relatively successful and may provide mechanistic insight into the origins of disease, it is also clear that variants can cause disease via other mechanisms such as removing key residues in an active site or perturbing interactions or regulatory mechanisms. Thus, methods used to predict the pathogenicity of missense variants often combine analysis of sequence conservation with information on protein structure and stability and other sources of information (***Kumar et al., 2009***; ***Adzhubei et al., 2010***; ***De Baets et al., 2012***; ***Kircher et al., 2014***; ***Choi and Chan, 2015***; ***Ioannidis et al., 2016***).

In the same way that perturbed protein stability may cause disease, protein structure, stability and folding also puts restraints of how amino acid sequences evolve (***Mirny and Shakhnovich, 1999***; ***DePristo et al., 2005***; ***Liberles et al., 2012***). Obviously, however, other considerations such as intrinsic activity and interactions with other molecules also play important roles in determining how sequences evolve, and site-to-site variation in evolutionary rates appear to be determined by a complex relationship between effects on stability and other functional constraints (***Echave et al., 2016***; ***Jimenez et al., 2018***; ***Echave, 2019***).

In order to understand better the relationship between protein stability, abundance and function we here asked the question of what fraction of single amino acid changes in a protein causes loss of function via loss of stability and cellular abundance of the proteins. Until recently, mutational analyses of proteins have mostly relied on a one-by-one approach in which individual amino acid changes are introduced and effects on various properties of a protein are tested—often using *in vitro* experiments on purified proteins (***Shoichet et al., 1995***; ***Fersht, 1999***). Such experiments can now be complemented by experiments that simultaneously probe the effects of thousands of variants in a single assay. Such *multiplexed assays of variant effects* (MAVEs, also often termed deep mutational scans) are based on developments in high-throughput DNA synthesis, functional assays and sequencing techniques (***Kinney and McCandlish, 2019***). Briefly, a selection procedure (e.g. for growth rate or a fluorescent reporter of a protein property) is applied to a large library of variants, each expressed in individual cells. Variants change in frequency depending on how they perform under the conditions of the selection, and the frequency of each variant before and after the selection is determined using next-generation DNA sequencing. Changes in variant frequency are used to compute a score that describes each variant’s effect on the property under selection. Such data can be used as an input to protein engineering (***Araya et al., 2012***; ***Shin and Cho, 2015***), to map local regions of fitness landscapes and help elucidate genotype-phenotype relationships (***Hietpas et al., 2011***; ***Sarkisyan et al., 2016***; ***Fernandez-de Cossio-Diaz et al., 2020***), and to understand which and how mutations may cause disease (***Starita et al., 2015***; ***Weile and Roth, 2018***; ***Stein et al., 2019***).

Now, for the first time, we have available measurements of thousands of variant effects on two key protein properties, activity and abundance, measured in multiple proteins. Here we take advantage of these data to examine more broadly how substitutions affect activity and stability. We examine how variants may affect abundance and activity differently to find functionally important positions in proteins (***Chiasson et al., 2020***), and to understand whether different types of effects are found in different regions of a protein’s structure.

To do so, we here analyse two different types of MAVEs that probe different aspects of protein function. As subjects of our study we have chosen two medically relevant human proteins, PTEN (phosphatase and tensin homolog) and NUDT15 (nucleoside diphosphate-linked to x hydrolase 15), because for both of these proteins multiplexed functional data exist from two different assays: One measuring the effect of variants on the activity of the protein via a growth rate (***Mighell et al., 2018***) or drug-sensitivity (***Suiter et al., 2020***) phenotype, and an assay that probes the effects of amino acid changes on cellular abundance (***Matreyek et al., 2018***; ***Suiter et al., 2020***). We will sometimes refer to the abundance data as reporting on ‘stability’ and the growth-based activity data as ‘activity’ or ‘function’, recognizing that the experiments report on a complex interplay of effects during the experimental assays. Notably, low scores in the activity-based assays might occur both due to loss of intrinsic enzymatic function, but also e.g. due to decreased protein abundance. Indeed, we use the complementary information on protein abundance to disentangle effects on abundance and intrinsic activity.

PTEN is a 403 amino-acid residue long lipid phosphatase expressed throughout the human body, and mutations in the PTEN gene have been associated with cancer and autism spectrum disorders (***Yehia et al., 2019***). In mice, PTEN has been shown to suppress tumor development via dephosphorylation of phosphatidylinositol lipids, although *in vitro* PTEN has been shown to have a broader range of substrates including proteins. PTEN is composed of two domains: a catalytic tensin-like domain (residues 14-185) and a C2 domain (residues 190-350) that mediates membrane recruitment (***Lee et al., 1999***). The C-terminal region of PTEN is disordered with a PDZ-domain binding region (residue 401-403) (***Valiente et al., 2005***). Our analysis of PTEN includes a MAVE that probes the effects of most single amino acid substitutions when assayed for lipid phosphatase activity in yeast (***Mighell et al., 2018***), whose growth had been made dependent on the ability of PTEN to catalyse the formation of essential phosphatidylinositol bisphosphate (PIP2) from its triphosphate (PIP3). While these experiments only probe one function of PTEN and might be affected also e.g. by expression levels, it has been shown that the resulting data accurately classifies the pathogenicity of PTEN variants (***Mighell et al., 2018***; ***Jepsen et al., 2020***). We complement these data with results from a different MAVE in which variant effects on cellular abundance are determined in an experiment termed ‘variant abundance by massively parallel sequencing’ (VAMP-seq) (***Matreyek et al., 2018***). In VAMP-seq the steady state abundance of protein variants in cultured mammalian cells is detected by fusion to a fluorescent protein, and cells are sorted using fluorescent activated cell sorting. The outcome of the VAMP-seq experiment is not substantially affected by the fusion with full-length GFP and correlates with *in vitro* measurements of thermal stability (***Matreyek et al., 2018***), but importantly also captures other effects that might affect protein abundance and which could be relevant for function, evolution and disease. Our analysis here covers the 56% of all possible single amino acid variants in PTEN for which we have measurements for both the activity and abundance, and thus complements our recent analysis of a small number of disease variants in PTEN (***Jepsen et al., 2020***).

NUDT15 is a nucleotide triphosphate diphosphatase that consists of 164 amino acids in a nudix hydrolase domain featuring a conserved nudix box that coordinates the catalytic Mg^2+^. The biologically relevant assembly is reported to be a homodimer although the monomer also has catalytic activity (***Carter et al., 2015***). NUDT15 deficiency is associated with intolerance to thiopurine drugs (***Yang et al., 2014***; ***Moriyama et al., 2016, 2017***; ***Nishii et al., 2018***), which are widely used in the treatments of leukemia and autoimmune diseases (***Karran and Attard, 2008***). Thiopurines are a class of anti-metabolite drugs that form the active metabolite, thio-dGTP, which competes with dGTP and causes apoptosis when incorporated extensively into DNA. NUDT15 hydrolyses thio-dGTP and thus negatively regulates the levels and cytotoxic effects of thiopurine metabolites. Therefore, NUDT15 variants that decrease function are a major cause of toxicity during thiopurine therapy, and thus the dose of the drug may be personalized to match the metabolism of these compounds (***Relling et al., 2019***). The high drug sensitivity of cells with compromised NUDT15 function has been used in a MAVE to assay 95% of all single amino acid variants for causing intolerance towards thiopurine drugs (***Suiter et al., 2020***). The same library and cells were also used in a VAMP-seq experiment to probe variant effects on cellular abundance, and like in the case of PTEN the results were shown to correlate with *in vitro* measurements of thermal stability. As in the case of PTEN, the outcome of the MAVE might depend on the exact conditions and e.g. drug concentration used, but was shown to capture the effects of several known pharmacogenetic variants (***Suiter et al., 2020***).

Here we have analysed the effect of variants on activity and cellular abundance in both PTEN and NUDT15 to provide a global view of what fraction of variants cause substantial loss of activity in the cell, and what fraction of these variants do so via loss of protein abundance. We find that approximately one third of all variants cause loss of protein activity, and that about half of these do so most likely because of loss of protein abundance. Variants that cause loss of abundance are often found inside the protein core, while variants that cause loss of activity without affecting abundance are often found in functionally important positions including those involved in catalysis or that interact with substrates. We also find that we can predict rather accurately the positions where substitutions generally give rise to decreased abundance and activity, whereas it remains difficult to quantitatively predict the effects of individual variants. Together, our results provide further insight into the link between thermodynamic stability and evolutionary conservation and experimental studies of different properties of fitness landscapes.

## Results and Discussion

### Global analysis of variant effects

We collected data from multiplexed assays reporting on both the activity and abundance of a total of 2822 variants in NUDT15 (***Suiter et al., 2020***) and 3927 variants in PTEN (***Matreyek et al., 2018***; ***Mighell et al., 2018***) (Fig. S1). Scripts to repeat our analyses are available online at github.com/KULL-Centre/papers/tree/master/2020/mave-analysis-cagiada-et-al. Two-dimensional histograms reveal that most variants have high scores in both assays, indicating wild type-like abundance and activity under the conditions of the cellular assays (Figs. 1A and B).

**Figure 1.**
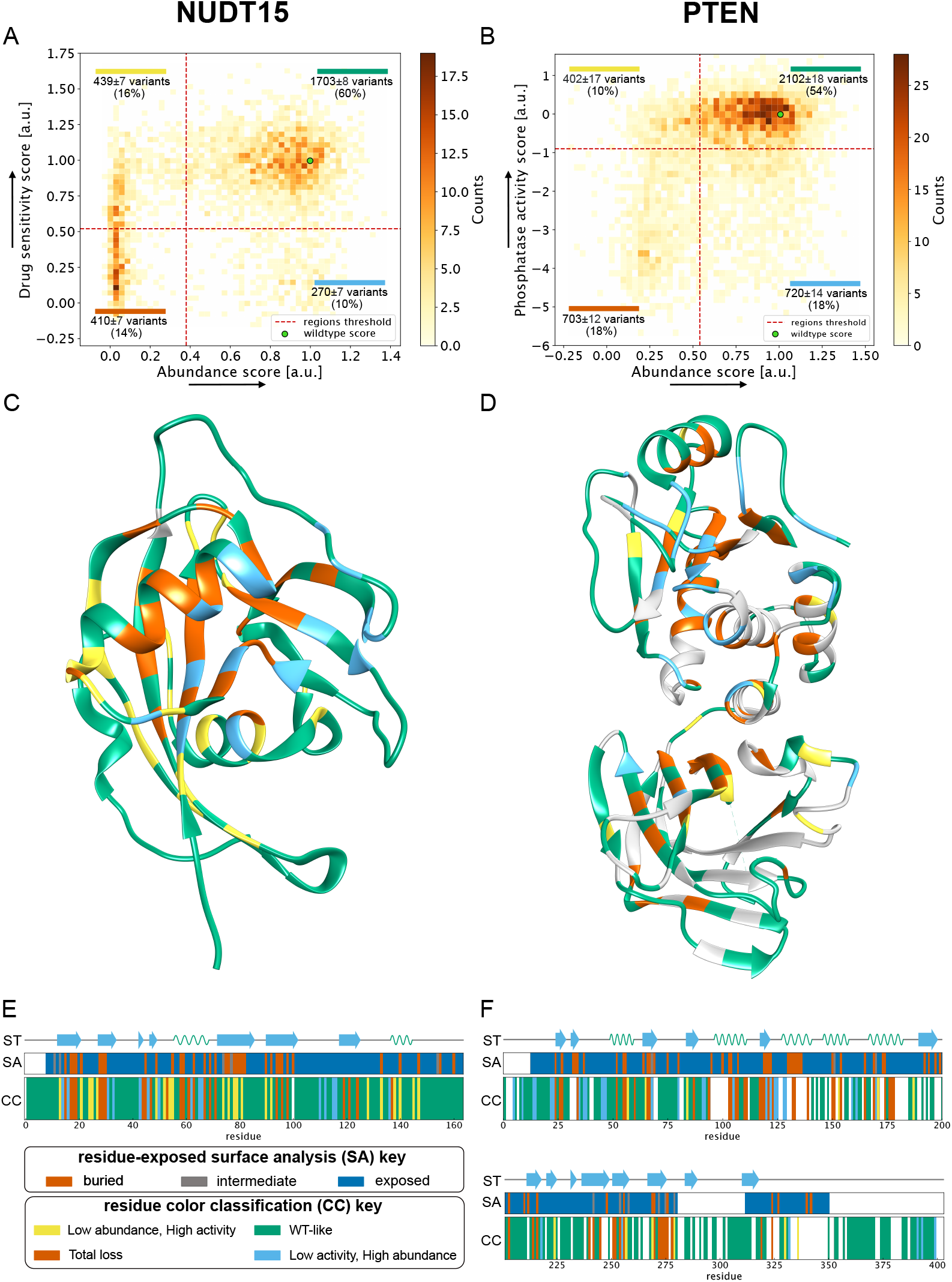
Overview of the NUDT15 and PTEN multiplexed data analysed in this work. A and B show 2D histograms that combine the data from the activity-based MAVE on the y-axis with the results from the VAMP-seq experiment on the x-axis. Variants are categorised based on the region of the 2D histogram (dashed lines) they belong to. The fractions of variants falling in each of the four quadrants are indicated, with errors of the mean estimated by bootstrapping using the uncertainties of the experimental scores. The two green points indicate the wild type. Arrows on the axes indicate direction of greater abundance or activity;for detailed definitions of the scores and their uncertainties we refer the reader to the original publications (***Matreyek et al., 2018***; ***Mighell et al., 2018***; ***Suiter et al., 2020***). Panels C and D show a per-position consensus category (CC) coloured onto the structure of the proteins (PDB entry 5LPG for NUDT15 and 1D5R for PTEN). Panels E and F show the positional colour categories together with the secondary structure (ST) and solvent accessibility (SA). The four classes of variants/positions are represented by a colour: ‘WT-like’ (green), ‘Low activity, high abundance’ (blue), ‘Low abundance, high activity’ (yellow) and ‘Total loss’ (red).

In order to separate wild-type like variants from those with decreased activity and/or abundance, we define a threshold value for all scores (Fig. S2). These thresholds define four classes of variants according to whether the variant showed high or low scores in the activity-based and abundance MAVEs. For simplicity each class is also associated with a colour. ‘WT-like’ variants had wild-type-like activity and abundance and are shown in green. ‘Low activity, high abundance’ variants had WT-like abundance but low activity in the assays, and are shown in blue. ‘Low abundance, high activity’ variants had WT-like activity but low abundance in the assays and are shown in yellow. ‘Total loss’ variants had low activity and low abundance and are shown in red.

For both proteins, the majority of variants are wild-type-like (60% for NUDT15 and 54% for PTEN; Figs. 1A and B; green). The total-loss category represents variants that both show loss of activity and low cellular abundance (14% for NUDT15 and 18% for PTEN; Figs. 1A and B; red), and as discussed further below we expect that most of these variants lose activity because of their low abundance. Of the total of 680 and 1403 variants with low activity in NUDT15 and PTEN respectively, 60% and 50% lose activity together with loss of abundance. The low activity, high abundance variants are still abundant in the cell but inactivated by other means, e.g. by changes in amino acids in the active site (Figs. 1A and B; blue). The low abundance, high activity class, which contains 16% of NUDT15 and 10% of PTEN variants (Figs. 1A and B; yellow), show low abundance levels, but high levels of activity in the activity-based assay and are not as easily explained by a single mechanism.

To focus our analysis on different types of variant effects in different parts of the protein structure, and to decrease uncertainty coming from examining individual variants, we converted the variant data into positional categories that represent the most frequent class among the variants at that position. We performed this classification procedure at all positions with at least five tested variants (99% for NUDT15 and 88% for PTEN), which also helped average out noise from examining individual variants with intermediate scores, and represent the classes using the same names and colouring scheme as for the variants. This results in 62% and 60% positions classified as WT-like for NUDT15 and PTEN respectively (Figs. 1C and D; green). On the other hand, at 15% and 22% of the positions most variants cause loss of activity together with loss of abundance (Figs. 1C and D; red), whereas loss of activity without loss of abundance is the most common outcome at 9% and 12% of the positions (Figs. 1C and D; blue). Finally, at 14% of the positions in NUDT15 and 6% in PTEN the variants most often have low abundance, but high levels of activity (Figs. 1C and D; yellow).

We validated the classifications using a clustering method that does not depend on defining cutoffs for the experimental scores. We grouped together positions with similar variant profiles in the two MAVEs (see Methods), and find overall very good agreement with the cutoff-based method in particular for the WT-like, total-loss and loss of activity, high abundance categories (Figs. S3 and S4). For NUDT15 we find that 133/163 positions are classified in the same way using the two different methods, with the most variable results occurring in the category with low abundance but sufficient activity to sustain growth (Fig. S3). For PTEN, we analysed the data using either three or four clusters, with the former appearing to be the more natural classification. In that case, 246/310 positions are classified in the same way using the two methods, with the 12 positions in the low abundance, high activity (yellow) category ending either as WT-like or total-loss. This indicates that three of the four categories of position effects are identified more robustly, corresponding to substitutions generally resulting in (i) WT-like activity in both assays, (ii) loss of activity and abundance or (iii) loss of activity, while retaining WT-like abundance. The low abundance, high activity positions are, however, less robustly classified and we do not analyse them further.

As expected, amino acids at buried positions are in general sensitive to mutations. In NUDT15, 35 out of the 163 amino acids are fully buried, and half of these (49%) are classified as sensitive to mutations in both the activity- and abundance-based assays (red label) with the remaining buried positions mainly classified as low abundance, high activity (34%; Figs. 1 E and F). Because the variant coverage is lower in PTEN, only 355 of 403 positions can be classified in this way, and only 34 of these 355 are fully buried. Among these 34, 80% are classified as ‘unstable’ positions (low abundance, high activity and total-loss categories). Thus, loss of abundance is the typical reason for loss of activity for variants at buried positions.

Low activity, high abundance positions are defined as having the majority of the tested variants that have lost activity, but are still abundant in the cell. Previously such positions have been found to map to functionally important sites in the membrane protein VKOR (***Chiasson et al., 2020***). We find that in PTEN, these variants and positions are mainly found in the catalytic phosphatase domain (Fig. S5) and include the active site (Figs. 2A and B). In NUDT15 we find the low activity, high abundance positions in several different regions. One group is located in proximity of the substrate binding site and include previously discussed Arg34 and Gly47 (***Suiter et al., 2020***). Another group includes the residues that coordinate a magnesium ion (***Suiter et al., 2020***). Finally, we find a group of residues that stretches from the substrate-interacting Arg34 and Gln44 (***Carter et al., 2015***) to Asn117 and Asn111 more distal from the substrate binding pocket and connected via a hydrogen bond network (Figs. 2C and D). Asn111 and Asn117 appear to help position a loop (residues 111-117) that includes the magnesium-coordinating Glu113, and although these residues do not directly contact the substrate, many substitutions lead to loss of function without loss of abundance.

**Figure 2.**
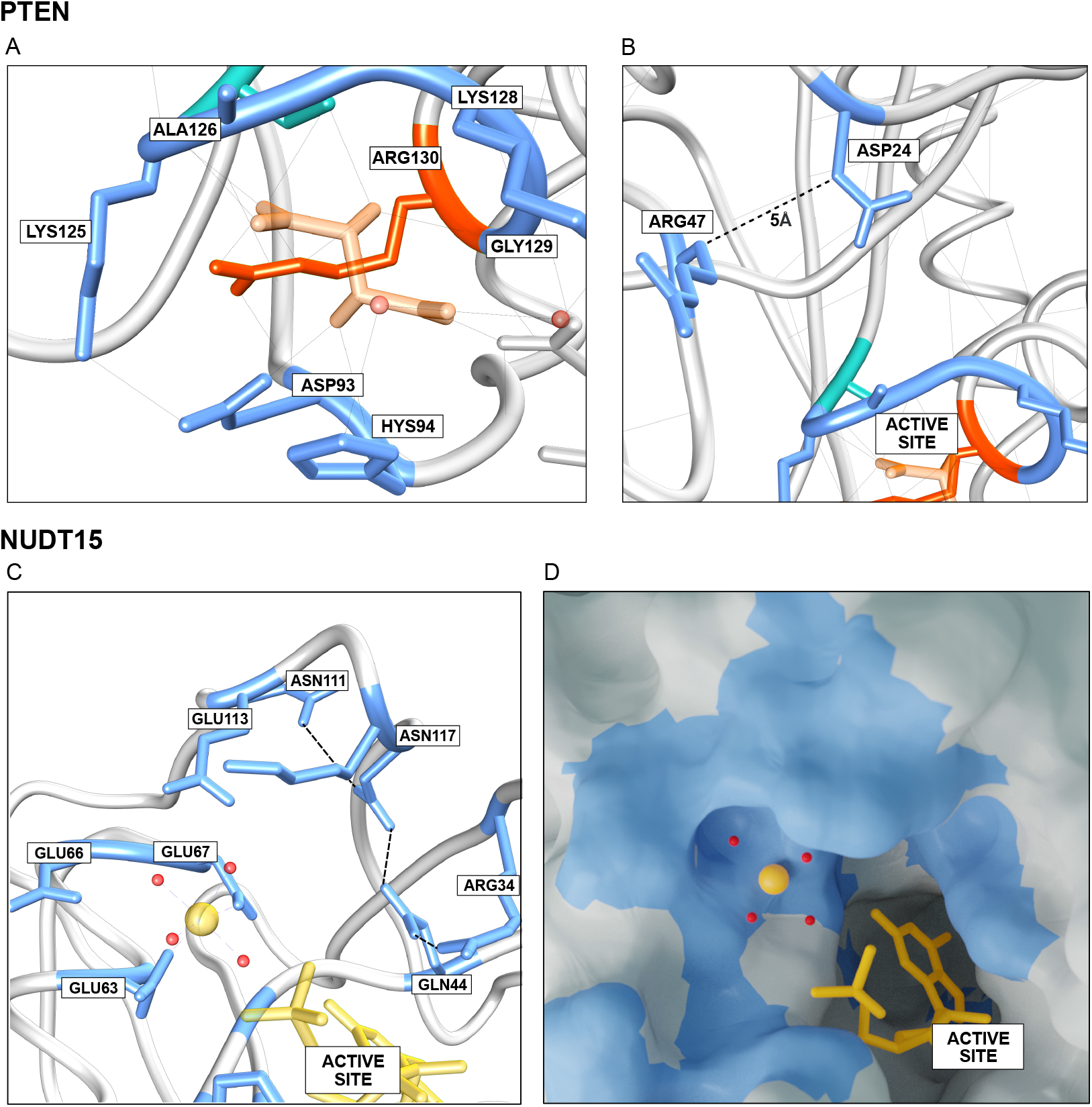
Examples of ‘low activity, high abundance’ positions. A: Residues in PTEN in the low activity, high abundance category (blue) include residues in and surrounding the catalytic phosphatase site including some that directly interact with the substrate (here mimicked by the inhibitor tartrate (***Lee et al., 1999***)). B: Other residues that are more distant to the active site also fall in this category, and variants in this region could perturb the integrity of the active site. C-D: Examples of functionally important residues in NUDT15 that are close to, but outside of the active site. In particular, we identified four conserved residues (Asn111, Asn117, Gln44, Arg44) that appear to connect via a hydrogen bond network, and whose perturbation could affect the hydrolysis of the thiopurines.

Having found that many low activity, high abundance positions play functional roles, we asked the question whether they are generally found near the active sites in NUDT15 and PTEN. Using Gly47 in NUDT15 and Arg130 in PTEN as reference points in the active sites in these two proteins we find that the low activity, high abundance positions, where variants typically show loss of activity, but not loss of abundance, are clustered around the active sites. Specifically, we find all of these positions in NUDT15 are within 14 Å of Gly47 (C*α*-distances). The average distance between low activity, high abundance positions and Gly47 is 9Å, a value that can be compared to the average (15Å) over all positions in NUDT15. In PTEN, we find that 29 of 32 low activity, high abundance positions are found in the catalytic domain. All of these 29 positions are within 22Å of Arg130, with the average distance to Arg130 being 14Å (compared to 21Å over all positions).

While the typical outcome at low activity, high abundance positions is loss of activity, not all substitutions have equally large effects, and some amino acid substitutions are more likely to be detrimental to function than others. We thus examined the individual substitutions at the low activity, high abundance positions positions to ask whether particular types of substitutions preserve function better than others. While such an analysis is made difficult by the small numbers of substitutions when broken down to start and end amino acid, we do find evidence suggesting differences depending on amino acid chemistry (Fig. S6). For example, at low activity, high abundance positions with Asn as the wild-type residue, it appears that substitutions to other small and polar amino acids (Asp, Ser, Thr) preserve function better than e.g. substitutions to hydrophobic amino acids (Fig. S6). More generally, it has previously been observed that there is a substantial effect of amino acid type on the outcome of a MAVE experiment (***Gray et al., 2017***; ***Dunham and Beltrao, 2020***), and we here find similar results (Fig. S7). As expected, we find that many total loss variants are substitutions of hydrophobic amino acids with charged and polar amino acids, while substitutions of hydrophobic residues with other hydrophobics are more common in the wild-type like category. More detailed analysis of these effects, however, are hampered by the low number of many of the types of substitutions.

### Computational predictions of multiplexed data from MAVEs

As described previously and demonstrated above, MAVEs provide a wealth of data not only for use in medical applications (***Weile and Roth, 2018***; ***Stein et al., 2019***) but also for understanding basic properties of proteins (***Dunham and Beltrao, 2020***). Despite recent advances in proteome-wide experiments (***Després et al., 2020***), it is still not possible to probe all possible variants in all proteins experimentally, and thus computational methods remain an important supplement to predict and understand variant effects. Experimental data from MAVEs are thus increasingly used to benchmark prediction methods, as they provide a broad view of the effect of amino acid substitutions in proteins (***Hopf et al., 2017***; ***Jepsen et al., 2020***; ***Livesey and Marsh, 2020***; ***Reeb et al., 2020***).

Recently we exploited the two different MAVEs for PTEN to analyse a small number of pathogenic variants together with variants that have been observed in a broader analysis of the human population (***Jepsen et al., 2020***). Specifically, we compared the abundance-based (VAMP-seq) and activity-based multiplexed data to two computational methods aimed at capturing either (i) specifically protein stability or (ii) function more broadly. Here we build on this work, by (i) applying computational modelling to predict changes in thermodynamic protein stability using Rosetta (***Park et al., 2016***) and (ii) using evolutionary conservation as a more general view of which amino acid changes would be tolerated while maintaining function (***Ekeberg et al., 2014***). The former uses as input the structure of NUDT15 or PTEN to predict the change in protein stability (ΔΔ*G*), while the latter uses a sequence alignment of homologuous proteins as input to a computational assessment of conservation, taking both site and pair-conservation (co-evolution) into account, quantified by a score (which we by analogy to ΔΔ*G* term ΔΔ*E*) that estimates how likely a substitution would be. We note that the same kind of model can be used to predict contacts in protein structure, but in line with previous work (***Lapedes et al., 2012***; ***Lui and Tiana, 2013***; ***Nielsen et al., 2017***; ***Hopf et al., 2017***) is here used to estimate the effects of amino acid substitutions. We also note that it has previously been shown that the ‘pair terms’ in these models, that capture effects of (apparent) co-evolution between pairs of sites, improve accuracy in these predictions (***Hopf et al., 2017***). As previously argued (***Jepsen et al., 2020***), the ΔΔ*G* calculations are more akin to the results of an abundance-based MAVE (both capturing aspects of protein stability), while the ΔΔ*E* values capture a broader range of effects as would also be expected from an activity-based MAVE.

We thus compared the computational predictions of ΔΔ*G* and ΔΔ*E* with each of the two multiplexed assays for NUDT15 and PTEN (Fig. S8). As expected, we find that stability predictions correlate better with the abundance-based MAVE than with the activity-based MAVE, while for the evolutionary analysis the situation is reversed (Fig. S9). In the case of NUDT15, for example, the data from the abundance-based MAVE correlates more strongly with the ΔΔ*G* calculations (*r_p_* = 0.57) than with ΔΔ*E* (*r_p_* = 0.42), whereas the activity-based MAVE is more poorly correlated with stability predictions (*r_p_* = 0.35) than with the conservation-based scores (*r_p_* = 0.52). While the difference is smaller (but still present) for PTEN, the results support the notion that analysis of conservation is a better predictor of general aspects of protein function, while the Rosetta calculations support the expected relationship between cellular protein abundance and thermodynamic stability (***Matreyek et al., 2018; Abildgaard et al., 2019***; ***Jepsen et al., 2020***). Also, we note that while the correlation coefficients are not very high, the results are in line with previous analyses of similar data (***Hopf et al., 2017***; ***Jepsen et al., 2020***; ***Livesey and Marsh, 2020***).

We define threshold values for the computational scores (Figs. S10 and S11) to separate wildtype-like from deleterious variants and construct four categories that we label with colours as above. Using a threshold of 2 kcal/mol for the ΔΔ*G* for both proteins results in 69% (NUDT15) and 65% (PTEN) of the variants being predicted stable. Similarly, from the evolutionary conservation analysis 78% and 58% of all variants for NUDT15 and PTEN, respectively, have scores that indicated that the substitutions are tolerated. Note that, by convention, positive ΔΔ*G* and ΔΔ*E* scores indicate loss of stability or sequence tolerance, respectively, and hence the scales are inverted compared to the scores from the MAVEs.

To enable a more direct comparison between the experimental and computational scores, we show histograms of the two computational scores (ΔΔ*G* and ΔΔ*E*) for each of the four classes based on the experimental scores (Fig. 3). We find that the variants that experimentally were classified as WT-like (stable and active) generally have low computational values; thus the computational predictions suggest that these substitutions have a mild effect on stability (low ΔΔ*G*) and are compatible with substitutions observed in homologous proteins (low ΔΔ***E***). We make similar observations for the total loss category, where the computational scores are generally above the cutoff, and for the low activity, high abundance category where the computational analysis finds low values of ΔΔ*G* but higher values of ΔΔ*E*. Despite these general trends, we find variable agreement in the classification of individual variants by experiments and computation (Fig. S12), with the best agreement in the WT-like and total-loss categories. To examine whether the results from the conservation analyses were specific to using lbsDCA, we also used the Evolution Trace algorithm (Lichtarge et al., 1996; Lua et al., 2016) to analyse the multiple sequence alignments, and found similar results (Fig. S13).

**Figure 3.**
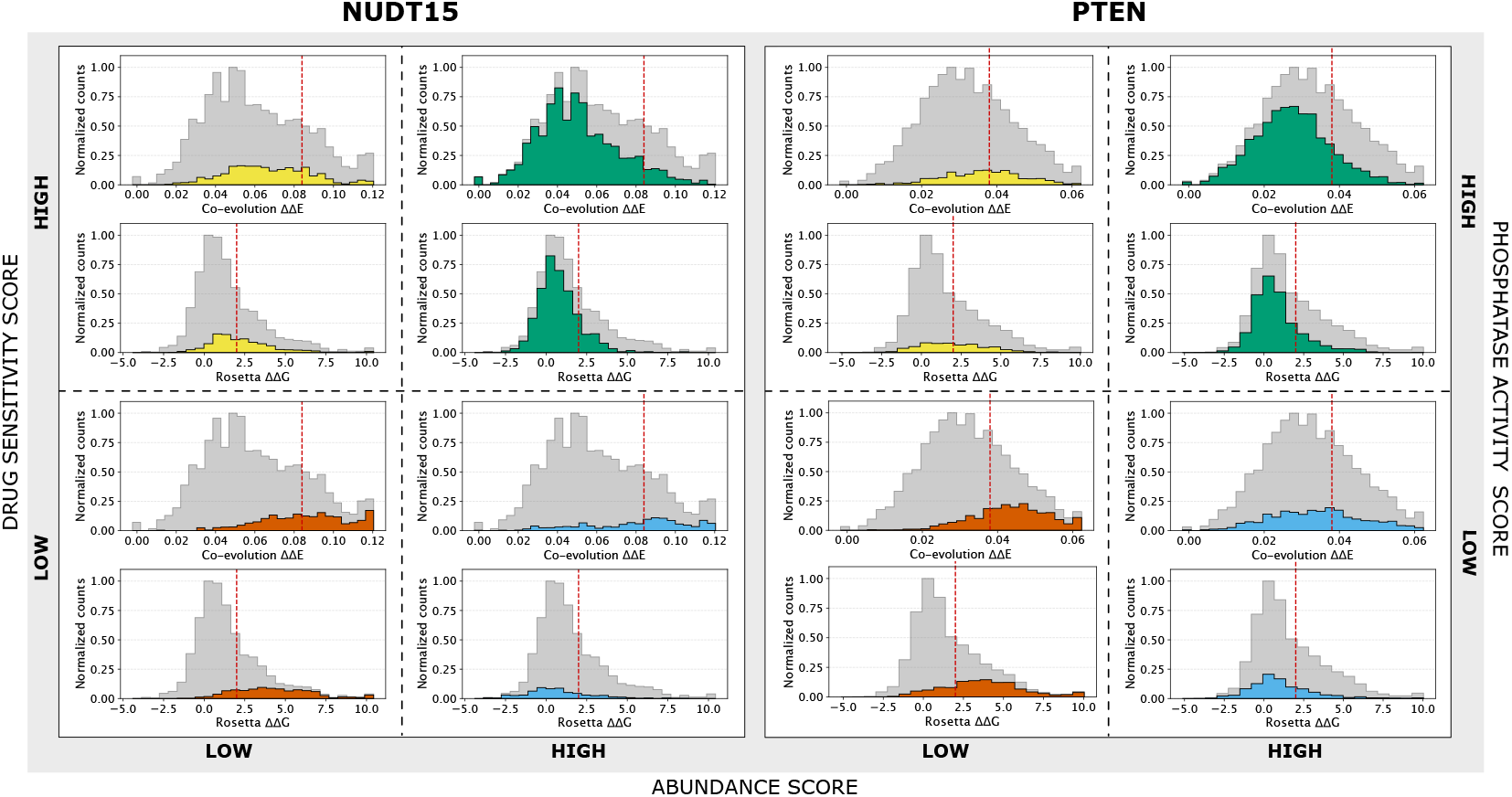
Histograms of the two computational scores (ΔΔ*G* and ΔΔ*E*) in NUDT15 and PTEN. ΔΔ*G* aims to capture effects purely on the thermodynamic stability, with high values indicating destablized variants. ΔΔ*E* captures evolutionary conservation, as calculated by a model that takes both site and pairwise co-evolution into account, and with high values indicating non-conservative substitutions. Thus, for both ΔΔ*G* and ΔΔ*E* positive values indicate detrimental substitutions, whereas in the experiments low values indicate substitutions that cause loss of activity or abundance. For both proteins we split the histograms up according to the four categories of variants determined from the experiments, as indicated by the axes with high and low experimental scores for abundance and activity. Thus, for example, the two green histograms for NUDT15 indicate the distributions of ΔΔ*G* and ΔΔ*E* values for those variants that are classified as stable and active by the MAVEs, and indeed it is clear that most of these variants have scores that are below the cutoff (red dashed lines). In addition to the coloured histograms we also show the full histogram of all analysed variants (grey) to ease comparison between the subsets and the full set of variants.

We proceeded by generating and examining the structure-function relationships that we extracted from the computational analyses (Fig. S14). We used the computational results to group the positions into four categories and found a substantial overlap with those found in experiments (Fig. S15), in particular for the WT-like and total-loss categories, with ca. 70% of the positions classified in the same way. This result suggests that the computational analyses better capture general effects at positions compared to individual variants as discussed above (Fig. 3). We again used a clustering procedure as an alternative approach to classify positions, and find good agreement both with the cutoff-based classification of the computational data as well as with the experiment-based classifications (Fig. S16). Thus, together these results show that a joint computational analysis of stability and conservation can be used to find positions in the protein where substitutions are likely to disrupt thermodynamic stability, and other positions where they will cause loss of activity via removing functionally important residues.

## Conclusions

Large-scale analysis of proteins using multiplexed assays provide opportunities to obtain a global view of variant effects (***Gray et al., 2017***; ***Dunham and Beltrao, 2020***). By combining different assays to read out different properties of a protein it becomes possible to dissect which positions contribute most to which property ***Jepsen et al., 2020***). Most proteins need to be folded to be active, and thus amino acid substitutions that lead to loss of stability will often lead to loss of function. Loss of stability thus appears to be an important driver for disease (***Yue et al., 2005***; ***Stein et al., 2019***) and determinant of evolutionary rates (***Echave et al., 2016***), and vice verse it has been shown that residues in active sites may be sub-optimal for stability (***Shoichet et al., 1995***).

We have here exploited the availability of data generated by MAVEs for two proteins, with one experiment probing general effects on protein activity and another directly assessing cellular abundance. We show that a global analysis of these experiments can provide insight into how proteins function and how activity may be perturbed. With the assays considered here, we find that most variants have at most a modest effect on protein activity. Of the ca. 30% of the variants that cause substantial loss of activity we find that ca. 50% also cause loss of abundance. Thus, while it is not surprising that there in many cases is a correlation between loss of function and loss of abundance, we here provide quantitative estimates on the relative importance of these effects across a wide range of substitutions in two unrelated proteins. The relative amounts of ‘low activity, high abundance’ and ‘total-loss’ variants that we find can be compared to our previous analysis of 42 disease-causing variants in PTEN, where we found a comparable fraction (~60%) of the disease-causing variants appears to cause loss of function via loss of stability and thereby cellular protein abundance ***Jepsen et al., 2020***). Indeed, most pathogenic variants in PTEN (***Mighell et al., 2018***; ***Jepsen et al., 2020***) score low in the activity based MAVE, with a substantial fraction of these also having low abundance (Fig. S17), while the situation for pharmacogenetic variants in NUDT15 (***Suiter et al., 2020***) is more complex (Fig. S17). Similarly, in our studies of pathogenic missense variants in the MLH1 gene we found low steady state protein levels (< 50% of wild type) in 7 out 16 pathogenic variants (***Abildgaard et al., 2019***). Thus, at least in these cases, it appears that the fraction of variants that cause disease via this mechanism reflects the overall fraction of ‘total loss’ variants in the protein. An interesting question for future experiments is how many of these variants would be active if protein levels could be restored for example by chemical chaperones or modulating the protein quality control apparatus (***Arlow et al., 2013***; ***Kampmeyer et al., 2017***). Indeed, chaperones are known to help buffer against destabilizing variants during evolution (***Rutherford and Lindquist, 1998***; ***Tokuriki and Tawfik, 2009***).

Building on previous work (***Cheng et al., 2005***; ***Chiasson et al., 2020***) we also show how we can use variant effects on protein activity and abundance/stability to find functionally important residues both by experiments and computation. For several surface-exposed residues many variants cause loss of activity, but without substantial loss of abundance. We find that these include the active sites in NUDT15 and PTEN, but also discover functionally important sites adjacent to these active sites. The importance of second shell positions for modulating the structure or dynamics of active site residues has for example also emerged in studies of ligand binding (***Tinberg et al., 2013***) and enzyme evolution and design (***Campbell et al., 2016***; ***Broom et al., 2020***). In our analysis of functional residues, we mostly focused on general effects at each position, rather than specific effects of individual substitutions. We did this to average out noise from individual measurements and to find general patterns, but with more data it would be interesting to perform such structure-sequence-function analysis at the level of individual substitutions.

The relatively tight confinement of these low-activity/high-abundance positions may also explain why predictions of changes in protein stability can be used to predict a substantial number of disease variants: At least in NUDT15 and PTEN the number of positions where substitutions typically cause loss of abundance (and thereby activity) is greater than the number of positions where substitutions cause loss of activity while retaining protein abundance. Indeed, while functional sites induce substantial constraints on amino acid variation during evolution, the strongest effects are those closest to the active sites (***Jack et al., 2016***; ***Mayorov et al., 2019***). Our ability to predict these sites by combining evolutionary analysis and stability calculations also suggest an approach for discovering new functionally-important sites using a combined analysis of protein structure and sequences. We find that ca. 12% of variants in NUDT15 and PTEN appear to be able to support wild-type like growth in the cellular assays even at substantially reduced protein levels. Clearly, there can be a non-linear relationship between a growth phenotype and protein abundance (***Jiang et al., 2013***), and this may help explain some of these variants. Future experiments that probe the relationship between expression levels and variant effects in NUDT15 and PTEN may shed further light on these variants. Further, the abundance-based MAVE for PTEN was performed in a cultured mammalian cell line (***Matreyek et al., 2018***) and the activity-based MAVE was performed in yeast (***Mighell et al., 2018***), leading to potential differences due to the differences in the quality control and proteostasis machinery in these cells.

In summary, we demonstrate how multiplexed assays and computational analyses are beginning to provide a coherent and comprehensive view of the global effects of variants in proteins. The results highlight that many effects are correctly predicted and thus computation can be used not only to predict whether a variant will cause loss of activity or not, but also provide some mechanistic insight. Clearly, there is room for improvement, and additional experiments on more proteins and covering more aspects of the complicated relationship between protein sequence and functions will help further our ability to predict these effects computationally (***Cheng et al., 2005***).

## Methods

### Conservation analysis of variant effects

We used a statistical analysis of multiple sequence alignments (MSAs) of the two proteins to estimate the tolerance towards specific substitutions. In line with previous work, we use a method that includes both site and pairwise conservation (co-evolution). We used the WT sequences from UniProt (P60484 and Q9NV35) as input to HHBlits (Remmert et al., 2012) to build initial MSAs, which we filtered before calculating the variant effects. The first filter removes sequences (rows) in the MSA with more than 50% gaps. The second filter keeps only positions (columns) that are present in the human target sequences of NUDT15 or PTEN. Finally, we apply a similarity filter (***Ekeberg et al., 2013***) to remove redundant sequences (more than 80% identical). We use a modified version of the lbsDCA algorithm (***Ekeberg et al., 2014***), based on *l*_2_-regularized maximization with pseudo-counts to predict the likelihood of every variant of the protein. We use the energy potential generated by the algorithm to evaluate the log-likelihood difference between the wild type and the variant sequences (ΔΔ*E*). We verified that the outcome of these analyses did not depend substantially on the parameters used to construct the MSA or to filter the alignments (Fig. S18). We performed Evolution Trace (***Lichtarge et al., 1996***; ***Lua et al., 2016***) calculations using the webserver available at evolution.lichtargelab.org.

### Structural analysis

We used Rosetta (GitHub SHA1 99d33ec59ce9fcecc5e4f3800c778a54afdf8504) to predict changes in thermodynamic stability (ΔΔ*G*) from the structure of NUDT15 and PTEN using the Cartesian ddG protocol (***Park et al., 2016***). As starting points we used the crystal structures of NUDT15 (***Valerie et al., 2016***) (PDB ID: 5LPG) and PTEN (***Lee et al., 1999***) (PDB ID: 1D5R). The values obtained from Rosetta were divided by 2.9 to bring them from Rosetta energy units onto a scale corresponding to kcal/mol (Frank DiMaio, University of Washington; personal correspondence) (***Jepsen et al., 2020***). We used DSSP-2.28 (***Kabsch and Sander, 1983***; ***Touw et al., 2015***) and the same crystal structures as above to classify the burial with a three state model (***Rost and Sander, 1994***) (buried, intermediate, or exposed).

### Defining thresholds for classifying variants

We defined thresholds for the scores from both MAVEs (Fig. S2), by fitting the variant score distributions using the minimal number of Gaussians (three) needed to obtain a reasonable fit. We then used the intersection of the first and last Gaussian as cutoff for our classifications. We use a cutoff of 2 kcal/mol (similar to the value used in our previous study (***Jepsen et al., 2020***)) for ΔΔ*G* and varied the cutoff for ΔΔ*E* to maximize the overlap in the classification of positions (Fig. S11).

To examine the threshold-based classifications, we used a hierarchical clustering algorithm (***Ward Jr, 1963***; ***Virtanen et al., 2020***) to group positions with similar responses to amino acid substitutions. Each position was represented by a 40-dimensional vector that contains the scores for each of the 20 possible amino acids in the two MAVEs. Missing values were replaced by the average score over that position. We use the Euclidean distance between these vectors as similarity score in the hierarchical clustering (***Ward Jr, 1963***). To compare with the threshold-based classification we analysed this using four clusters, though in the case of PTEN we also show the results using only three clusters.

### Residue classification

We assigned a category to residues for which data for at least five variants are available in both MAVEs. We used the mode (the most common class of the variants at that position) to assign the residue category.

## Data Availability

Code and data to repeat our analyses are available online at github.com/KULL-Centre/papers/tree/master/2020/mave-analysis-cagiada-et-al.

## Acknowledgement

This work is a contribution from the PRISM (Protein Interactions and Stability in Medicine and Genomics) centre funded by the Novo Nordisk Foundation (to R.H.-P., D.M.F., A.S. and K.L.-L.). J.J.Y. acknowledges funding from the NIH (R01GM118578).

## Supporting Material

**Supporting Figure S1.**
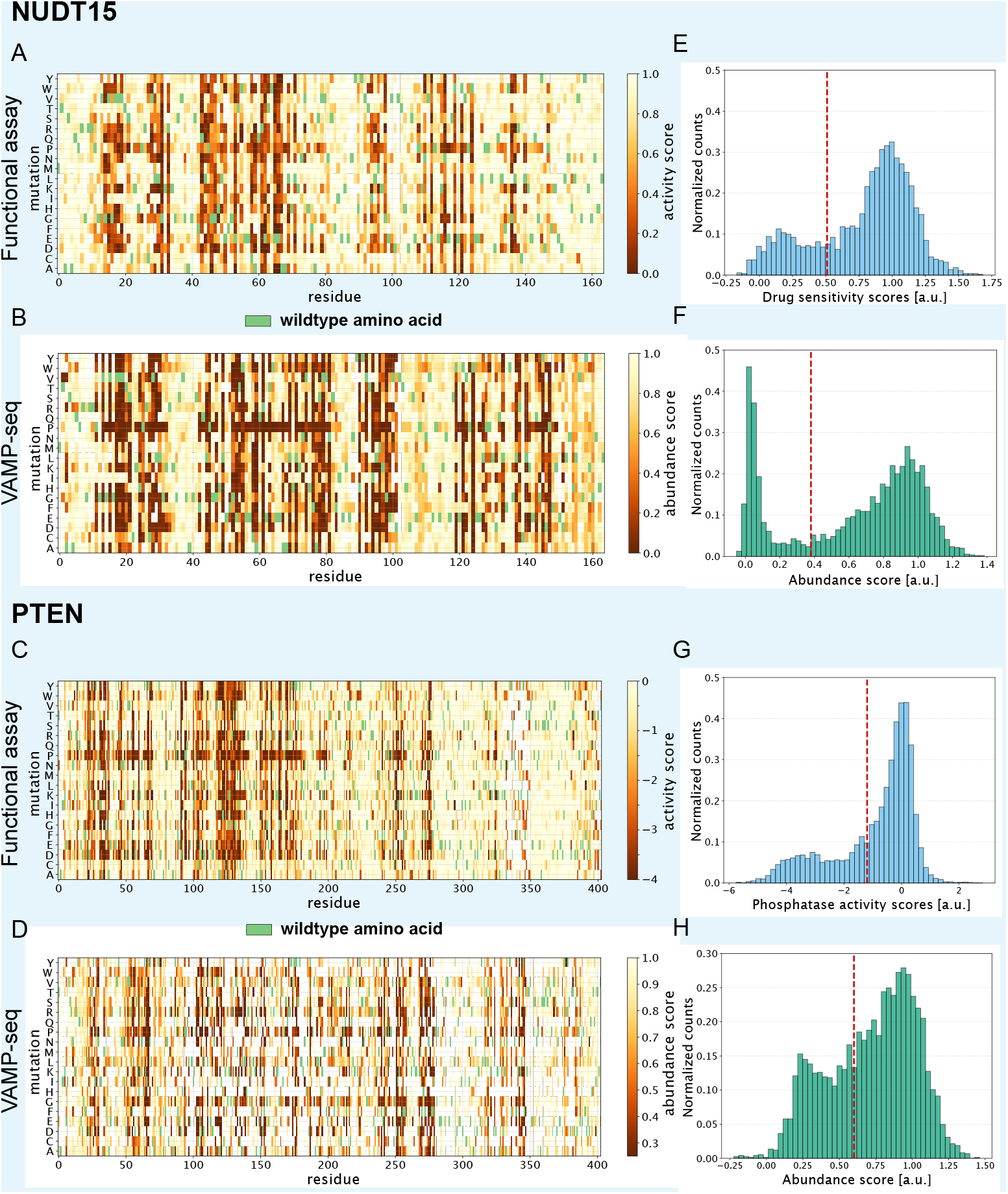
Experimental data for NUDT15 and PTEN. Panels A–D show the multiplexed data as heat maps. The experimental scores are indicated by colours and the wild type sequence is represented by a light green block at each position. Variants with wild-type-like behaviour have higher scores, and variants with lower scores indicate those of either low abundance (VAMP-seq) or activity (activity-based assays). Panels E–H show the distribution of values in each of the four assays with the red dashed lines indicating the cutoffs used to separate low and high scoring variants.

**Supporting Figure S2.**
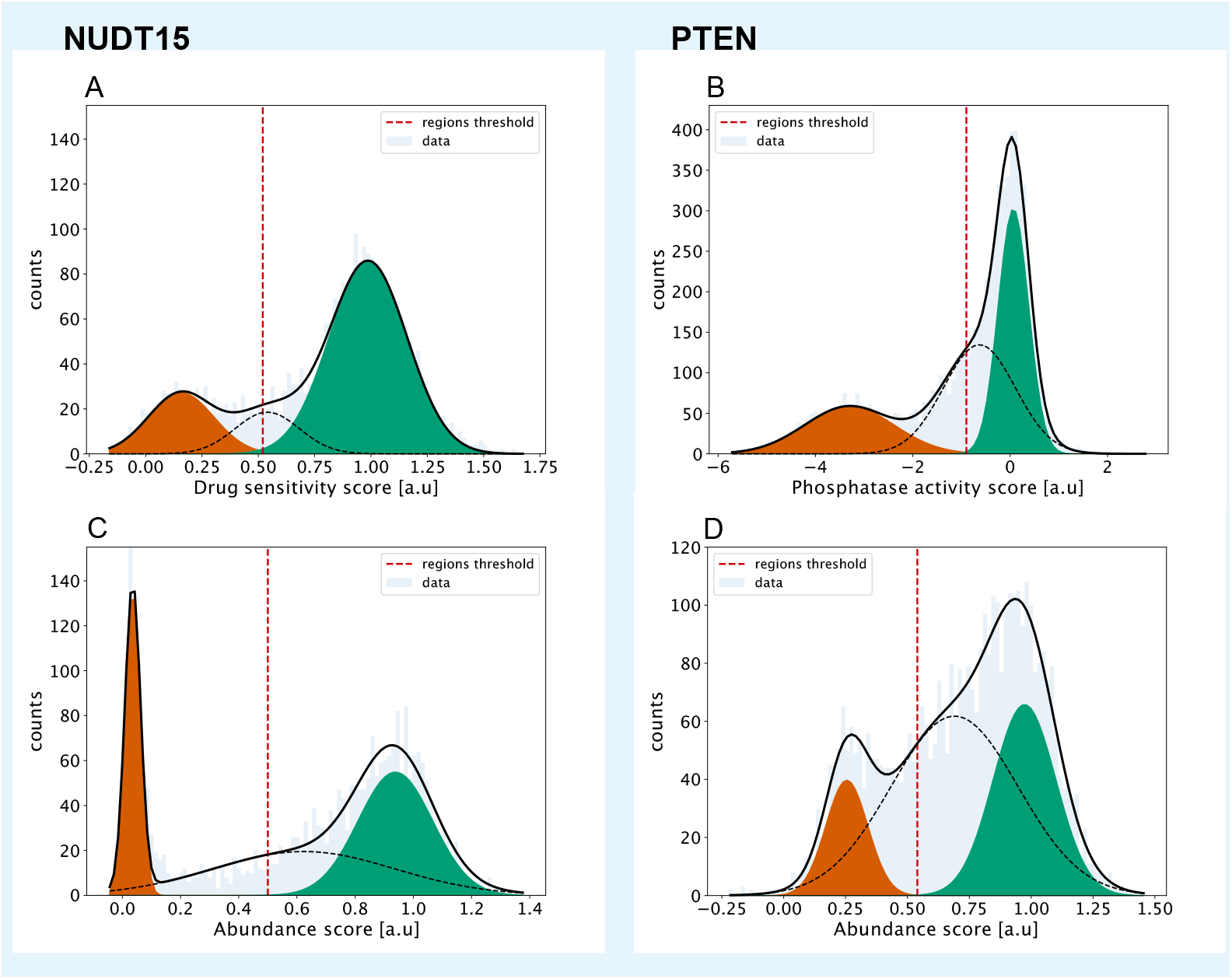
Protocol used to define thresholds for the multiplexed data generated by MAVEs. In each panel we fit the distribution of scores (light blue) to a mixture model with three Gaussian distributions (black line). The first and last Gaussian are shown in red and green, respectively, and we used the intersection between these to define cutoffs for classifying the variants (red dotted line).

**Supporting Figure S3.**
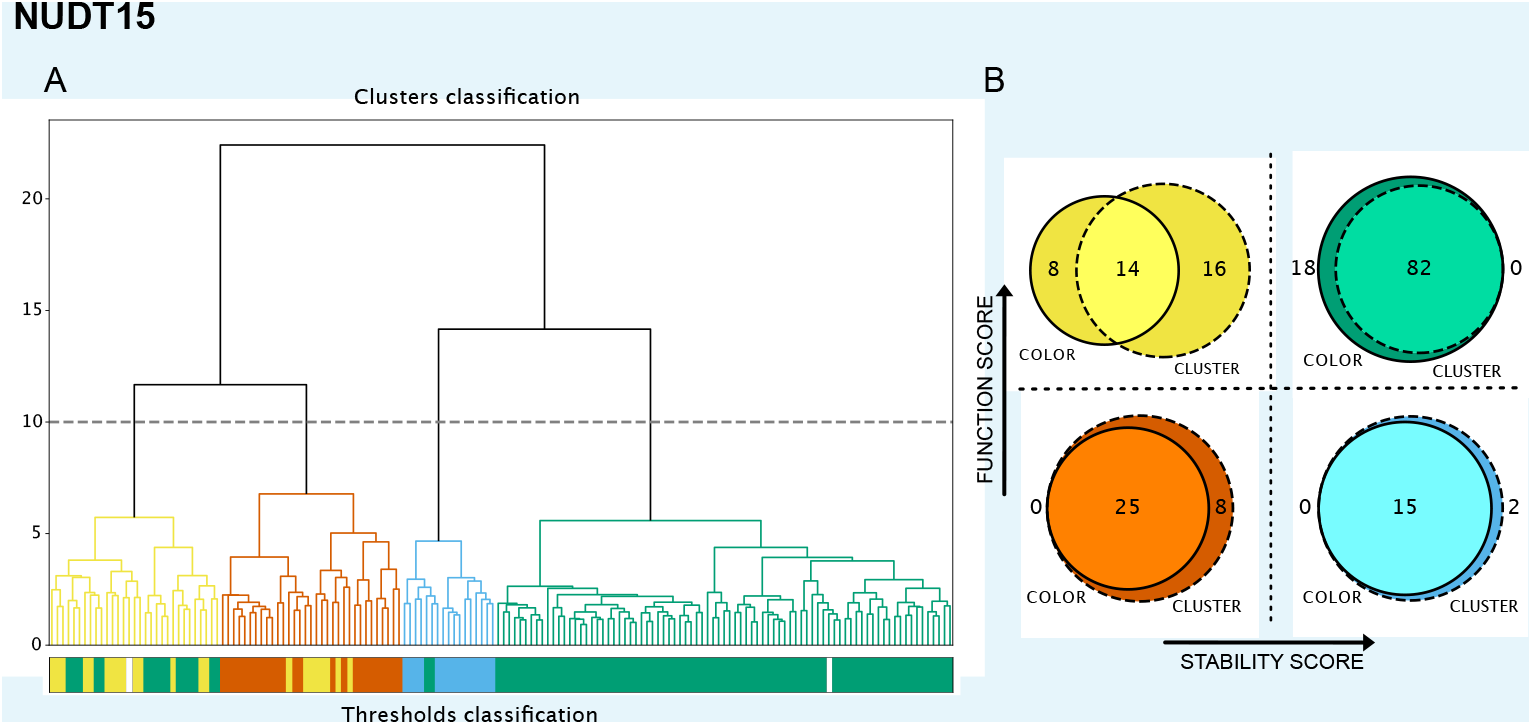
Assessment of the quality of a cutoff-based classification of positions in NUDT15. A: A hierarchical cluster analysis grouping together positions with similar responses to amino acid substitutions. Using the horizontal line to define the number of clusters, we colour the four clusters analogous to the results of the threshold-based classification. The bar plot under the cluster figure shows the colour assigned to each position in the threshold-based classification. B: Agreement between the two classifications represented using Venn diagrams.

**Supporting Figure S4.**
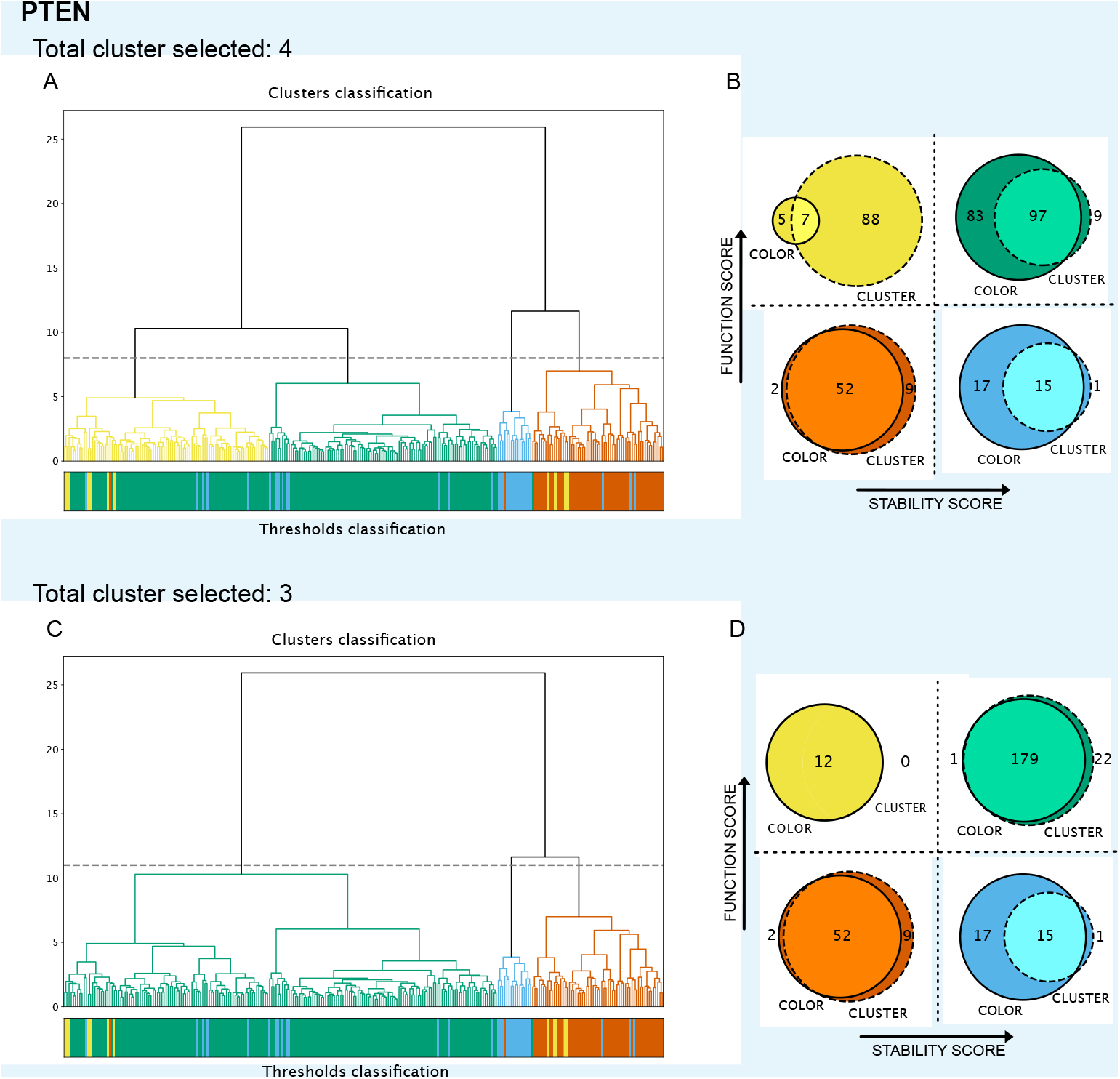
Assessment of the quality of a cutoff-based classification of positions in PTEN. A: A hierarchical cluster analysis grouping together positions with similar responses to amino acid substitutions. Using the horizontal line to define the number of clusters, we colour the four clusters analogous to the results of the threshold-based classification. The bar plot under the cluster figure shows the colour assigned to each position in the threshold-based classification. B: Agreement between the two classifications represented using Venn diagrams. Panels C and D use the same clustering as in A, but with the cutoff set so as to obtain only three clusters. In this case, the yellow group disappears and is merged with the green cluster.

**Supporting Figure S5.**
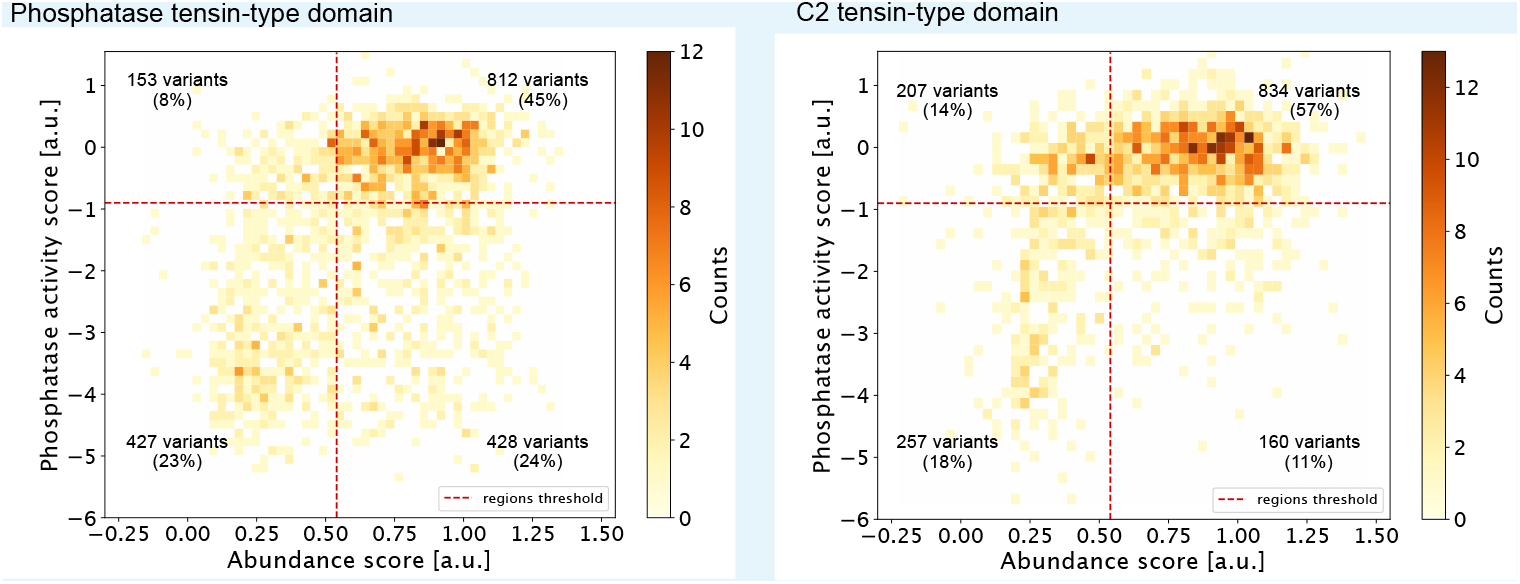
Analysis of PTEN variants by structural domain. The figures show the 2D histograms of the MAVE scores in PTEN for each of the two structural domains. While the overall picture in the two domains is similar, it is clear that a greater fraction of variants (47%) in the catalytic phosphatase domains causes loss of activity compared to the C2 domain (29%).

**Supporting Figure S6.**
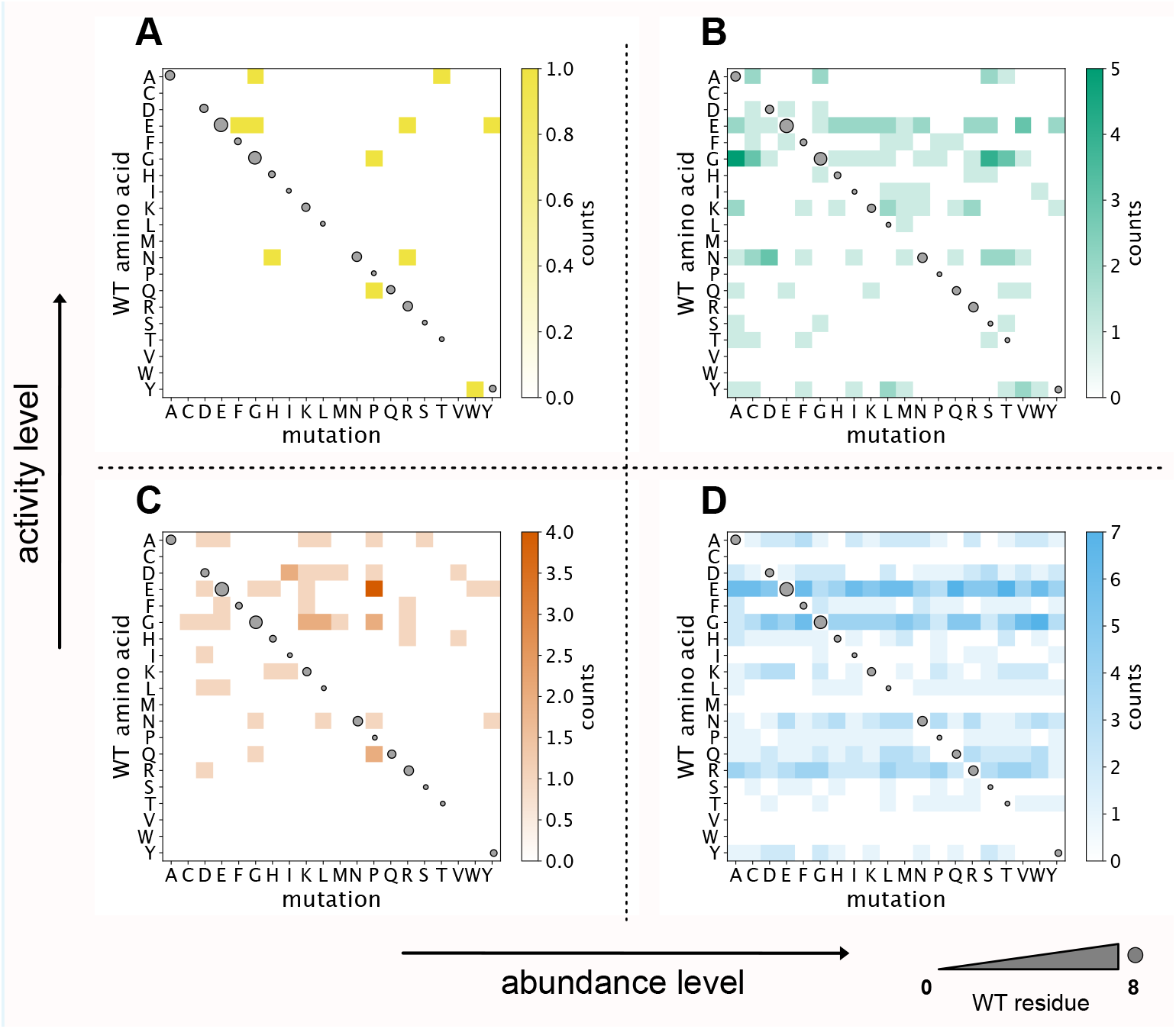
Statistics of types of amino acid substitutions at ‘low activity, high abundance’ positions in both NUDT15 and PTEN. For each type of substitution (pairs of ‘start’ and ‘end’ amino acid type) with data from the two MAVEs we calculated how many of these fall into each of the four categories. Because this analysis is focused only on those positions where the typical outcome is low activity, high abundance, there are most variants in this category. However, individual variants at these positions might behave differently, as indicated by the number of variants in the three other categories.

**Supporting Figure S7.**
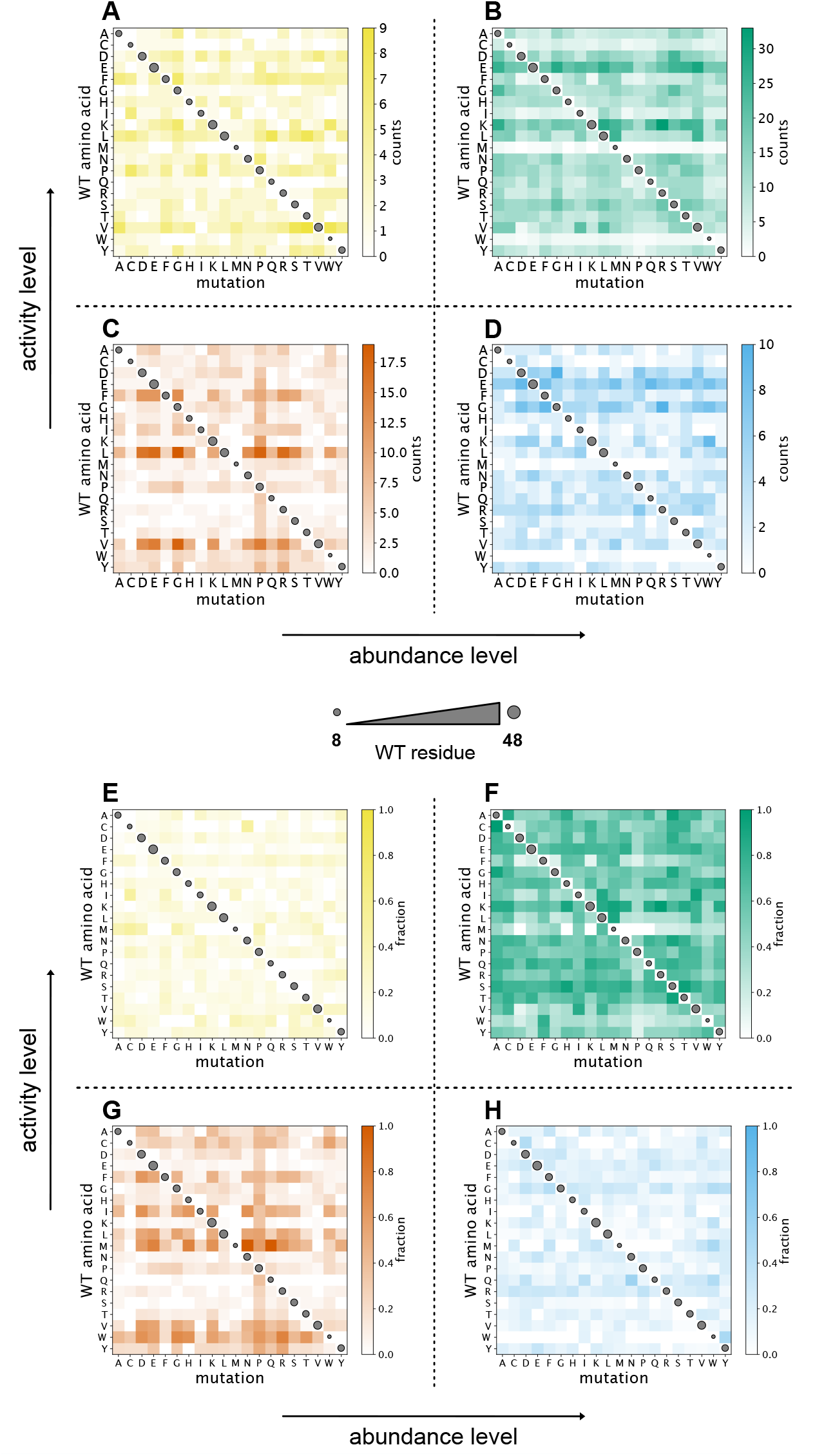
Statistics of types of amino acid substitutions over all positions in both NUDT15 and PTEN. For each type of substitution (pairs of ‘start’ and ‘end’ amino acid type) with data from the two MAVEs we calculated how many of these fall into each of the four variant categories. Panels A–D show the raw counts, whereas panels E–H are normalized so that the sum for a given pair of start/end amino acids is one across the four categories.

**Supporting Figure S8.**
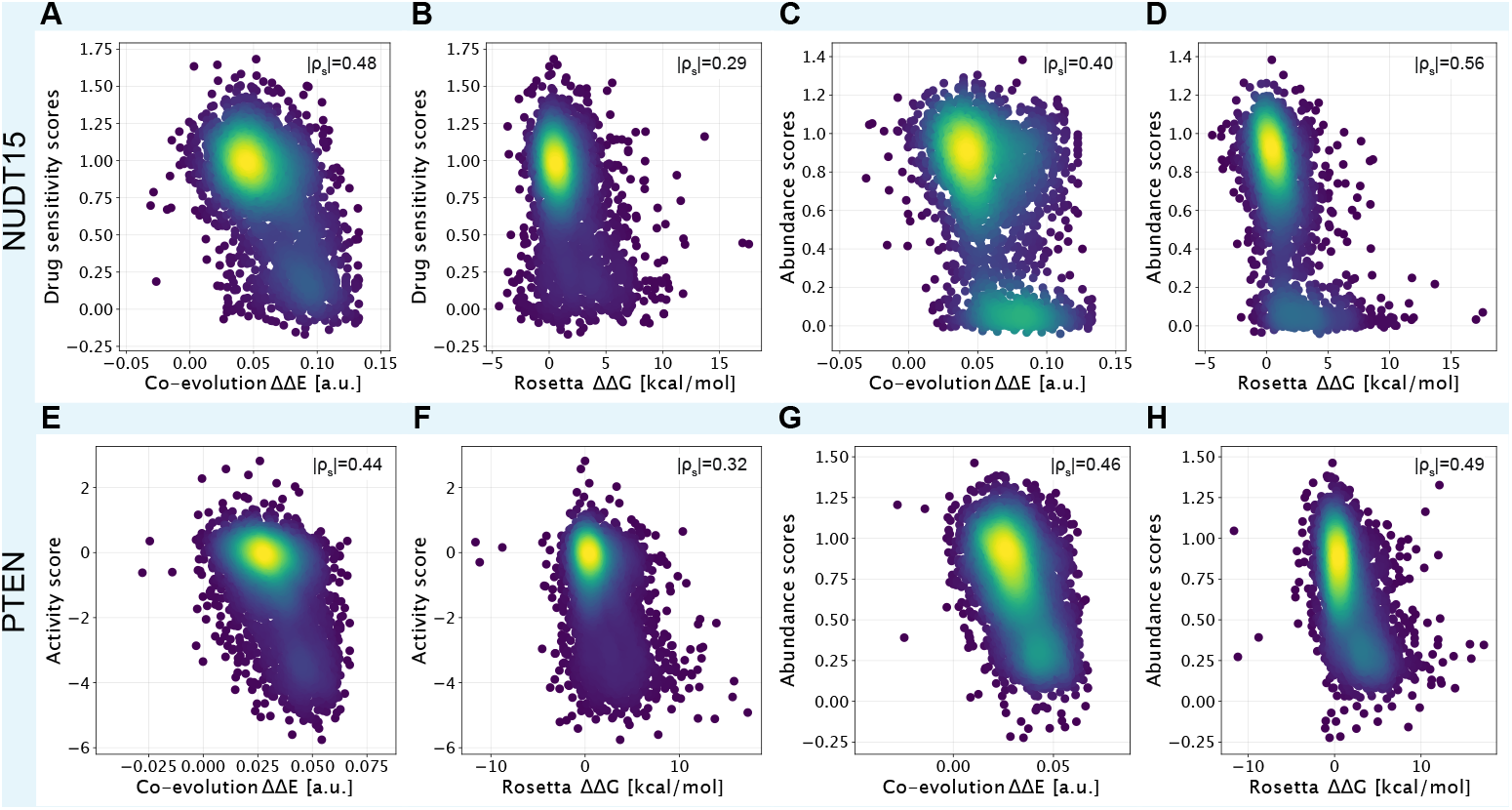
Score density scatter plots of data from MAVEs and computation. In each panel the computational scores are on x-axis and the MAVE scores on the y-axis. The different variant density is represented through the color scale with blue indicating low density regions and yellow high density regions. Panels A and E contain comparison between ΔΔ*E* and the activity-based MAVEs, and panels C and G between ΔΔ*E* and VAMP-seq. Panels B and F show the correlation between ΔΔ*G* and the activity-based MAVEs, and panels D and H between ΔΔ*G* and VAMP-seq.

**Supporting Figure S9.**
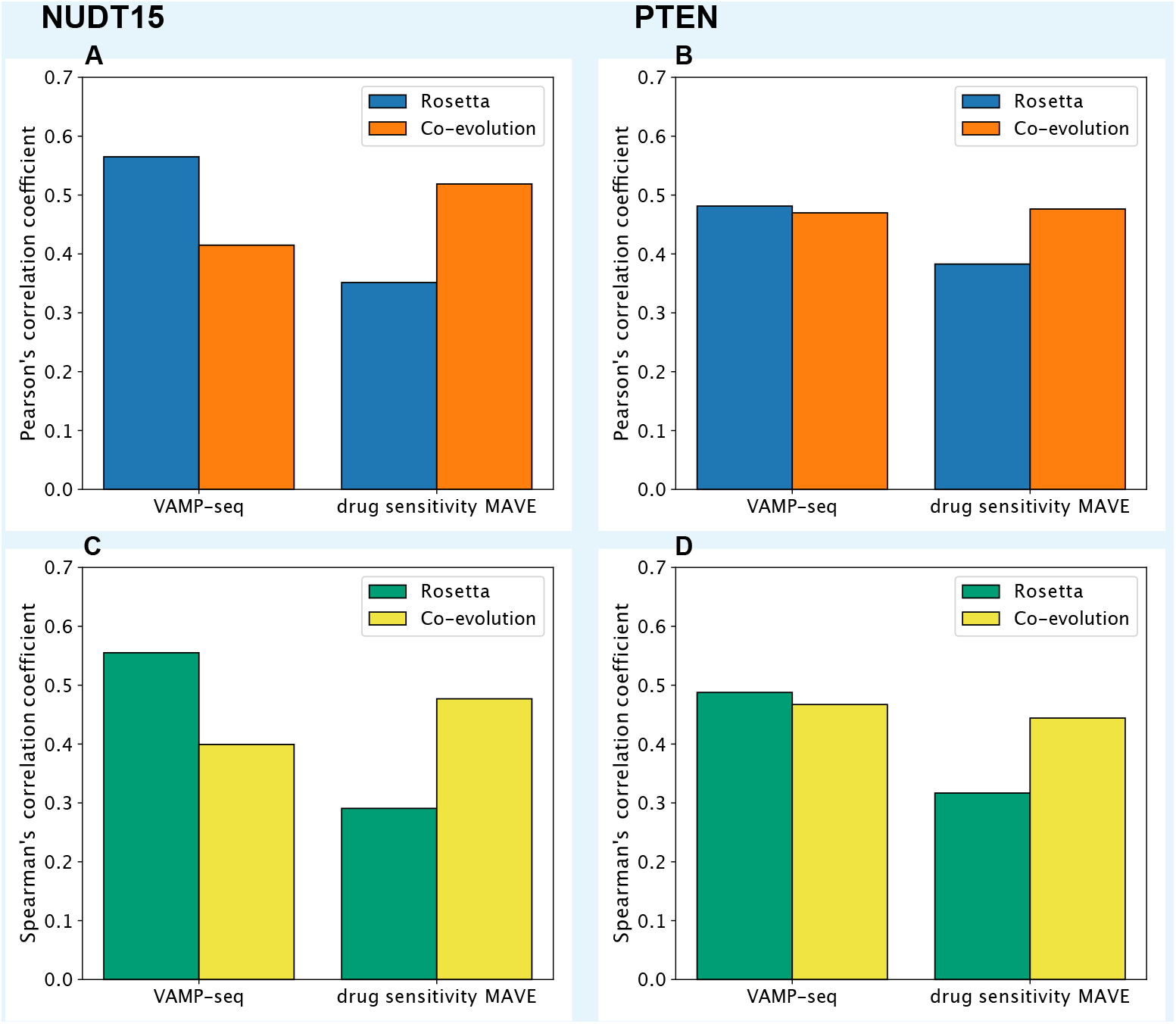
Correlation between the experimental data and two computational scores.In panels A and B we show, for each MAVE, the Pearson correlation coefficient to either (blue bars) the Rosetta stability predictions or (orange bars) an assessment of tolerance towards substitutions using an evolutionary model (co-evolution). Panels C and D show the same comparisons, but using Spearman’s correlation coefficient.

**Supporting Figure S10.**
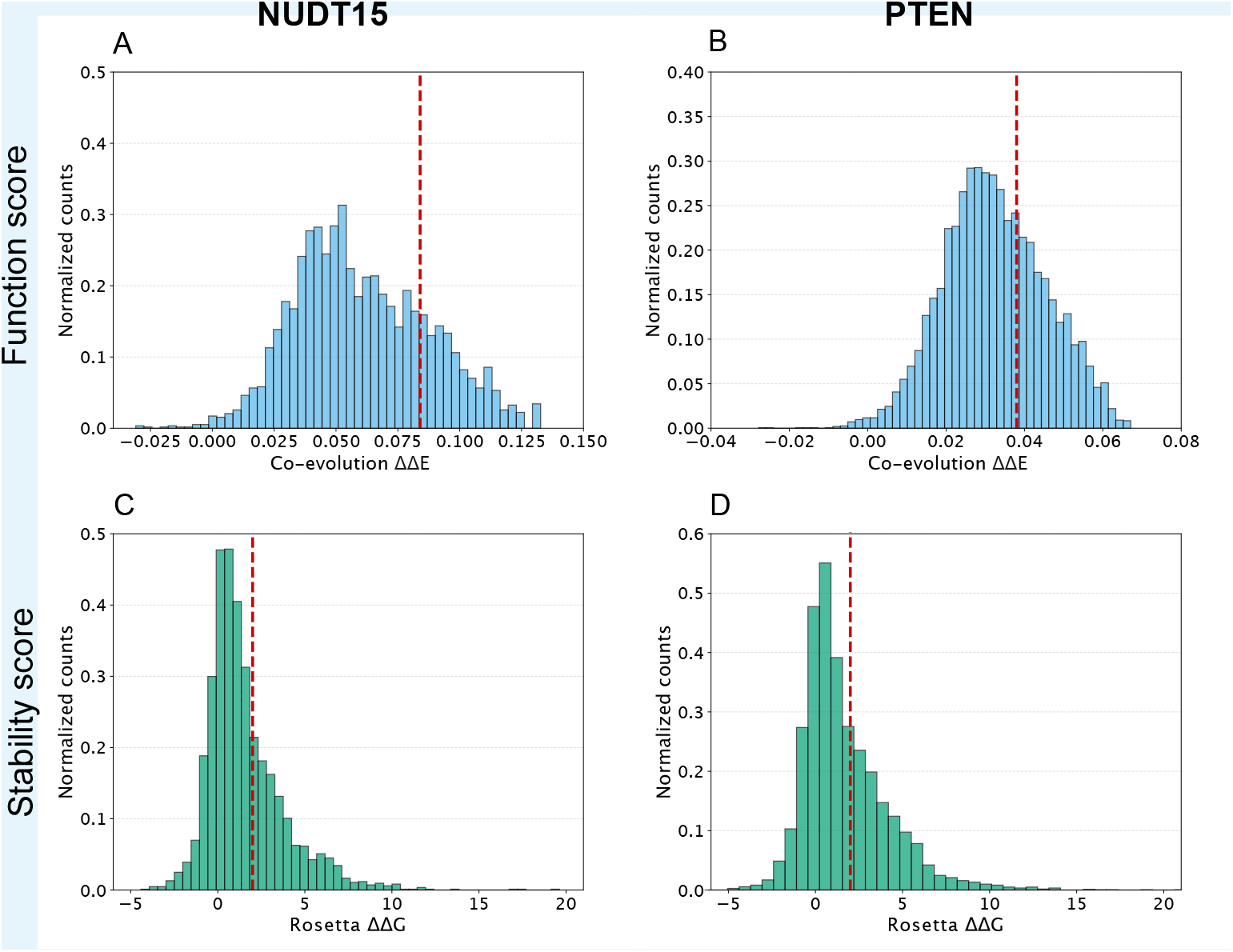
Distributions of scores and thresholds used in the computational analyses. Panels A and B show the distribution of values in the co-evolution analysis for both the target proteins, with the red dashed lines indicating the cutoffs used for classifying the variants. The Panels C and D show the distributions and the thresholds for the Rosetta ΔΔ*G* values.

**Supporting Figure S11.**
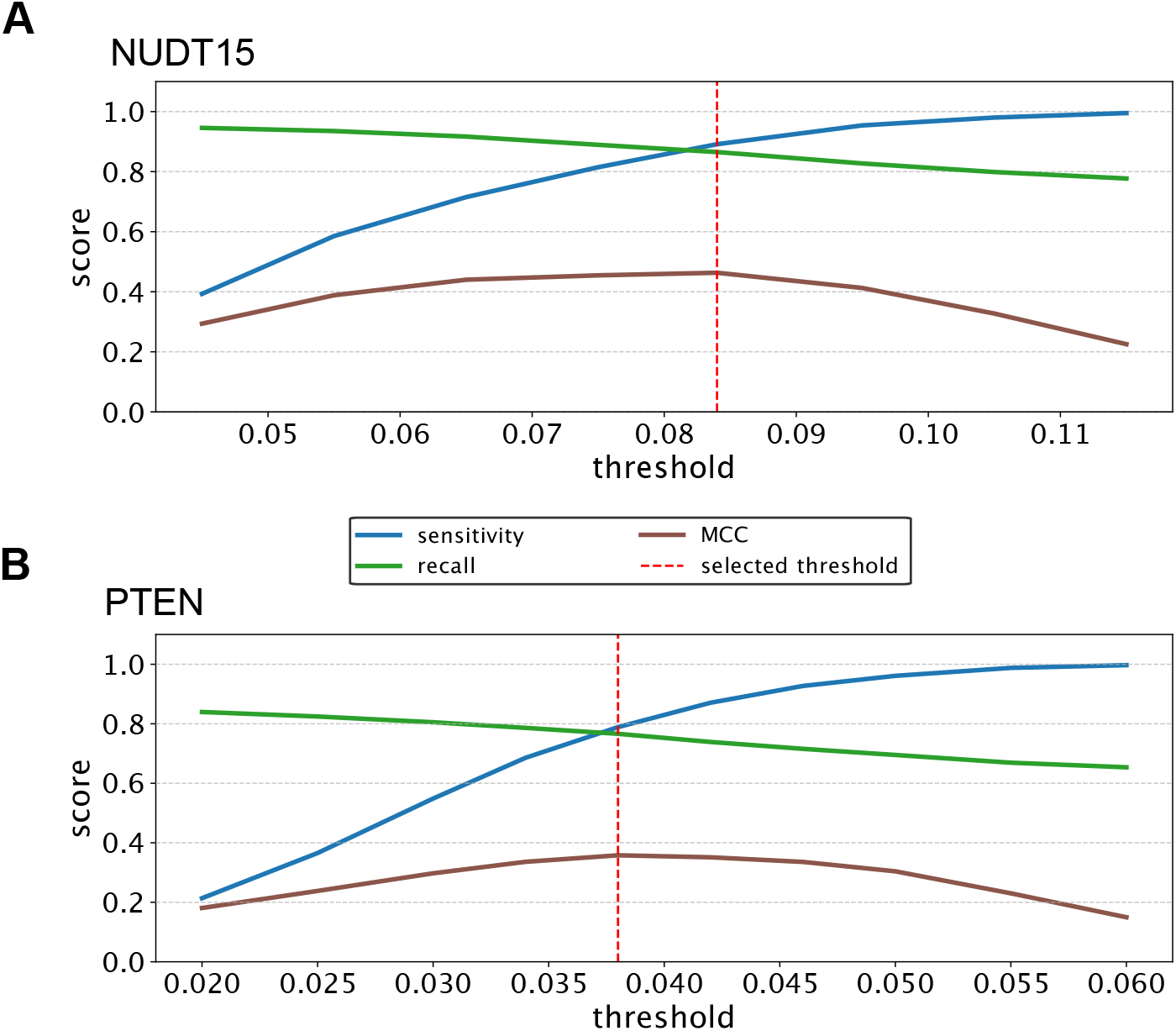
Selection thresholds for lbsDCA analysis of NUDT15 and PTEN. For both (A) NUDT15 and (B) PTEN we varied the cutoff used to separate conservative (low ΔΔ*E*) and non-conservative (high ΔΔ*E*) substitutions (see also Fig. S10). For each value we compared the results using the binary classification of low/high values in the two activity-based MAVEs (Fig. S2) and calculated the sensitivity, recall and Matthews Correlation Coefficient (MCC). The red lines show the values used in the remainder of the analysis.

**Supporting Figure S12.**
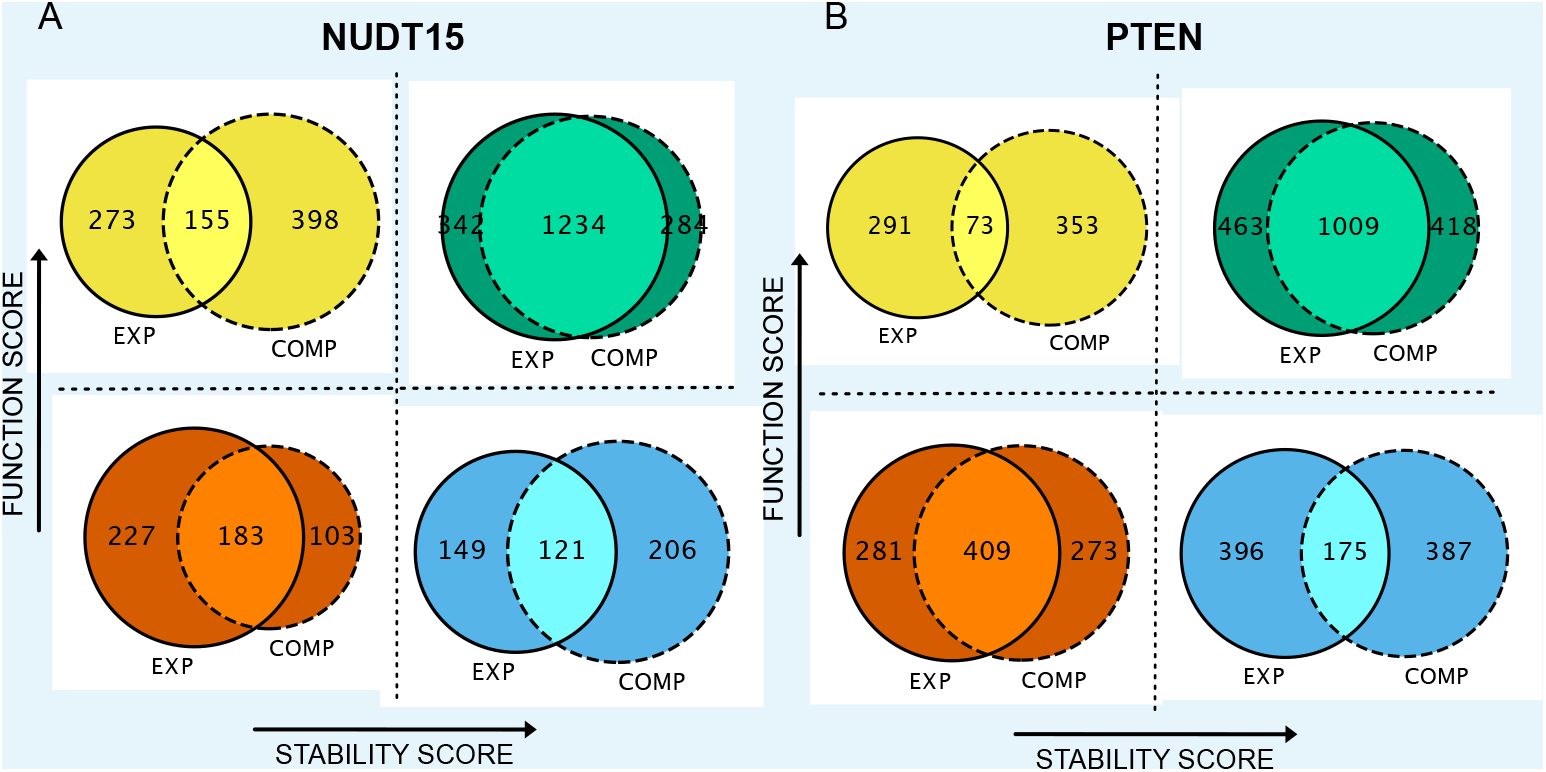
Comparison of the classification of variants by analysing the computational and experimental data analysis. The Venn diagrams show the agreement between the classification of the individual variants in (A) NUDT15 and (B) PTEN with ‘EXP’ and ‘COMP’ representing the cutoff-based classification using either the data generated by the two MAVEs or the ΔΔ*G*/ΔΔ*E* analysis, respectively.

**Supporting Figure S13.**
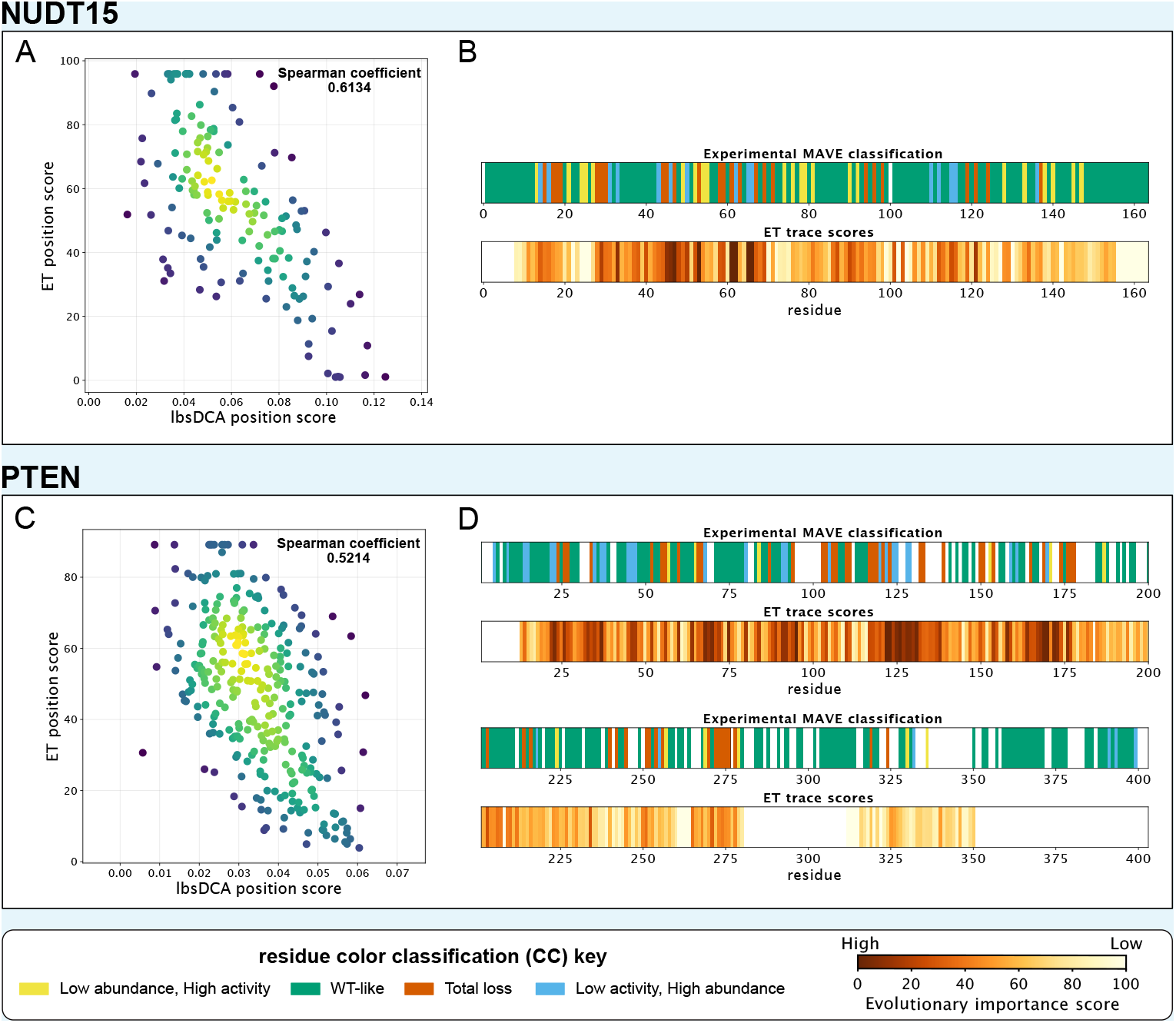
Sequence analyses using the Evolution Trace (ET) algorithm. Panels A and C show the correlation between the scores from ET and lbsDCA, with the latter calculated as the average over variant scores at each position. Panels B and D show a comparison of the MAVE-based classification of positions with the ET scores. A high value in ET analysis corresponds to a position that is conserved in evolution.

**Supporting Figure S14.**
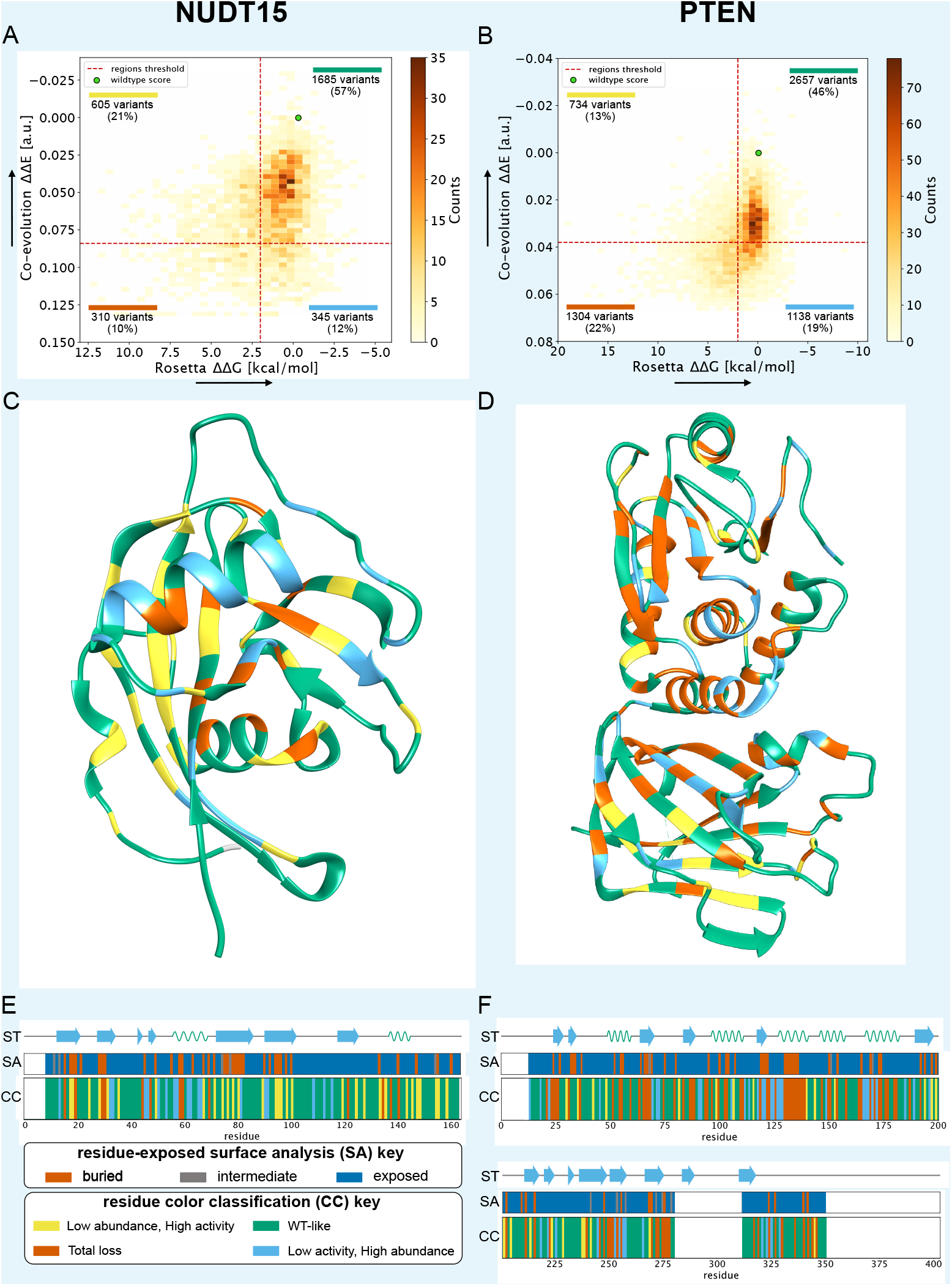
Overview of the NUDT15 and PTEN computational data generated in this work. Panels A and B show 2D histograms that combine the evolutionary conservation analysis ΔΔ*E* on the y-axis with the Rosetta ΔΔ*G* values on the x-axis. Variants are categorised based on the region of the 2D histogram they belong to. Green dot indicates the wild type. Panels C and D show a per-position consensus category (CC) coloured onto the structure of the proteins (PDB entry 5LPG for NUDT15 and 1D5R for PTEN). Panels E and F show the positional consensus categories together with the secondary structure (ST) and solvent accessibility (SA).

**Supporting Figure S15.**
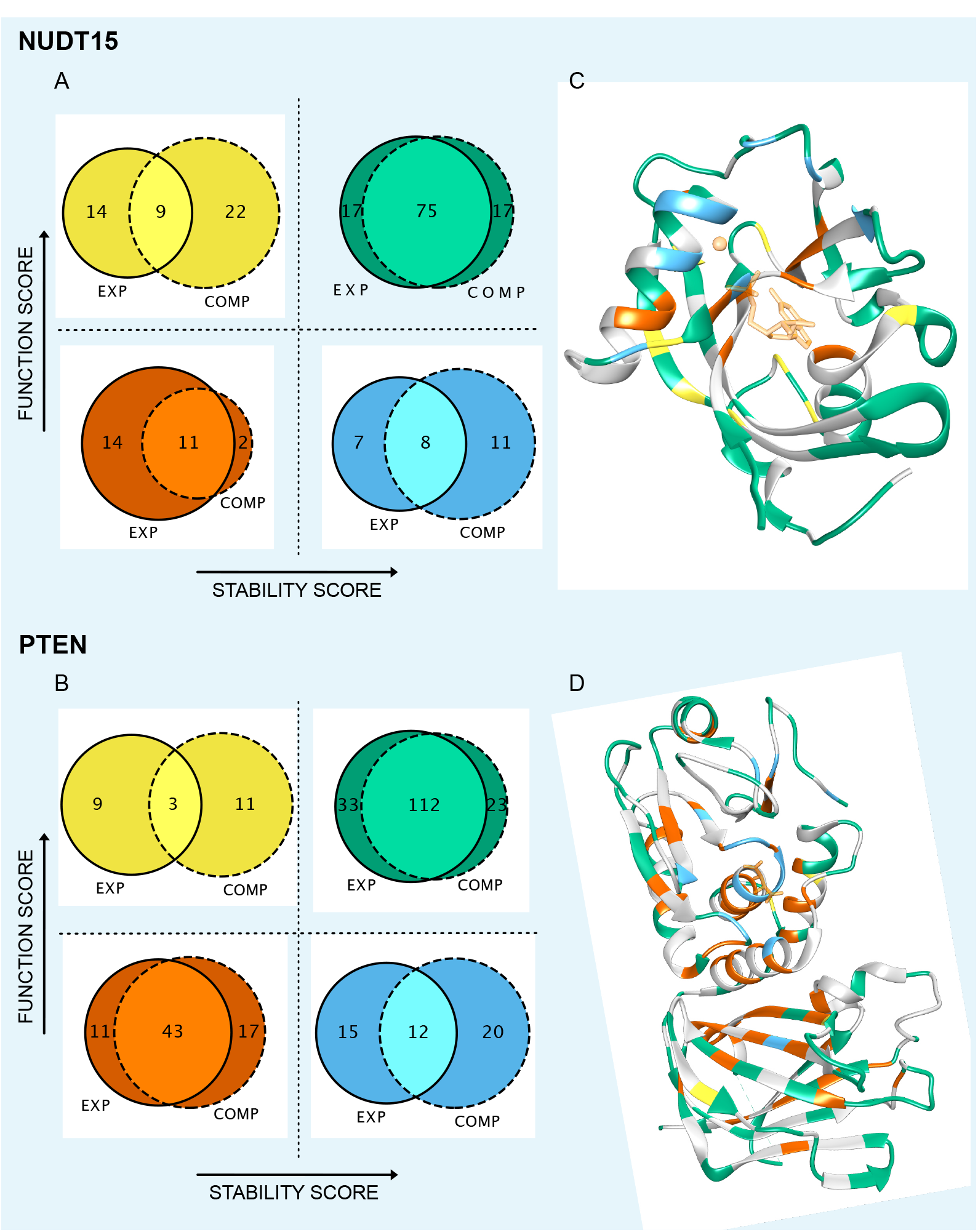
Comparing the classification of positions obtained using experimental data and the computational analysis. Panels A and B show the agreement between the classification from the analysis of experiments and computation (on the common subset of tested positions) using Venn diagrams. The protein structures in panels C and D are coloured at those positions where the experimental and computational classification is the same with the remaining positions shown as grey.

**Supporting Figure S16.**
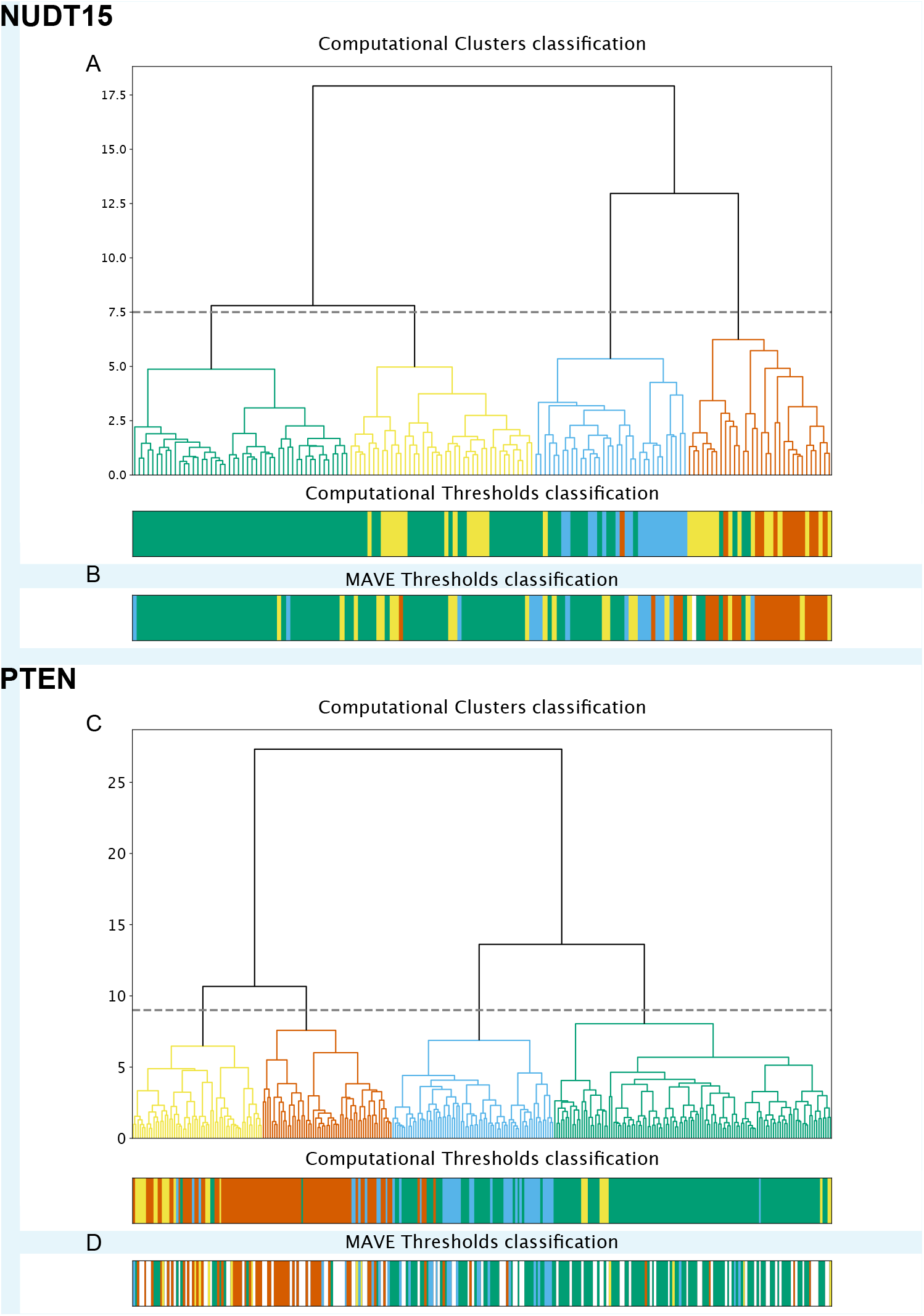
Cluster analysis of computational ΔΔ*E* and ΔΔ*G* data. The panels show the result of a hierarchical clustering of (A) NUDT15 and (C) PTEN. The clusters are shown with the same colours used to define the classes of position in the cutoff-based classification. The two bar plots (B and D) show the class assigned to each position in the cutoff-based classification using either computational or experimental data.

**Supporting Figure S17.**
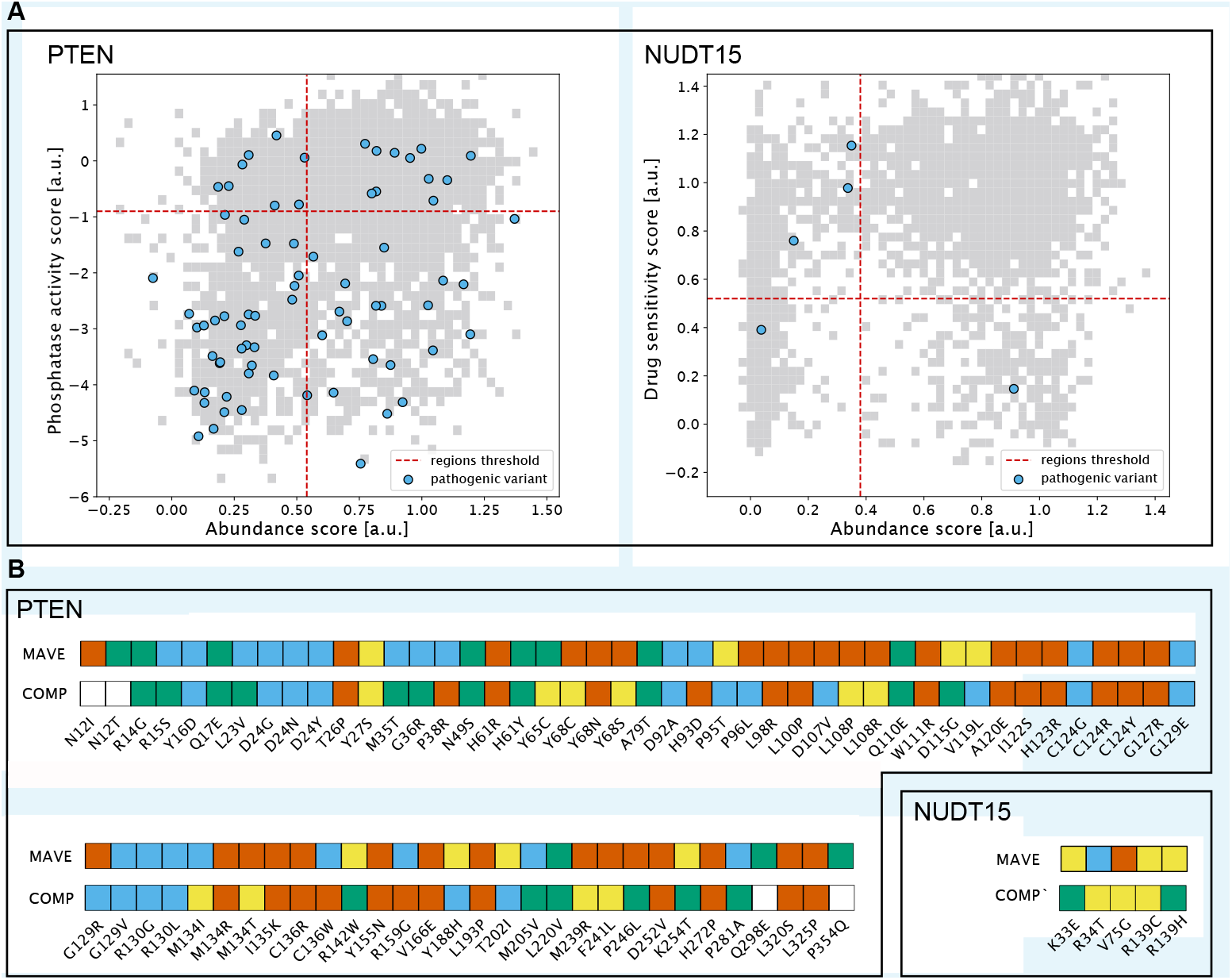
Analysis of known pathogenic variants in PTEN and pharmacogenetic variants in NUDT15. Panel A shows the MAVE scores for pathogenic variants (PTEN) and pharmacogenetic variants (NUDT15) (blue points) on a background of the full set of variants (grey). Panel B details the variant classification for these variants using both the MAVEs and computational analyses, using the same colour scheme as in the remaining text. Variants that are labelled white could not be classified in the computational analysis.

**Supporting Figure S18.**
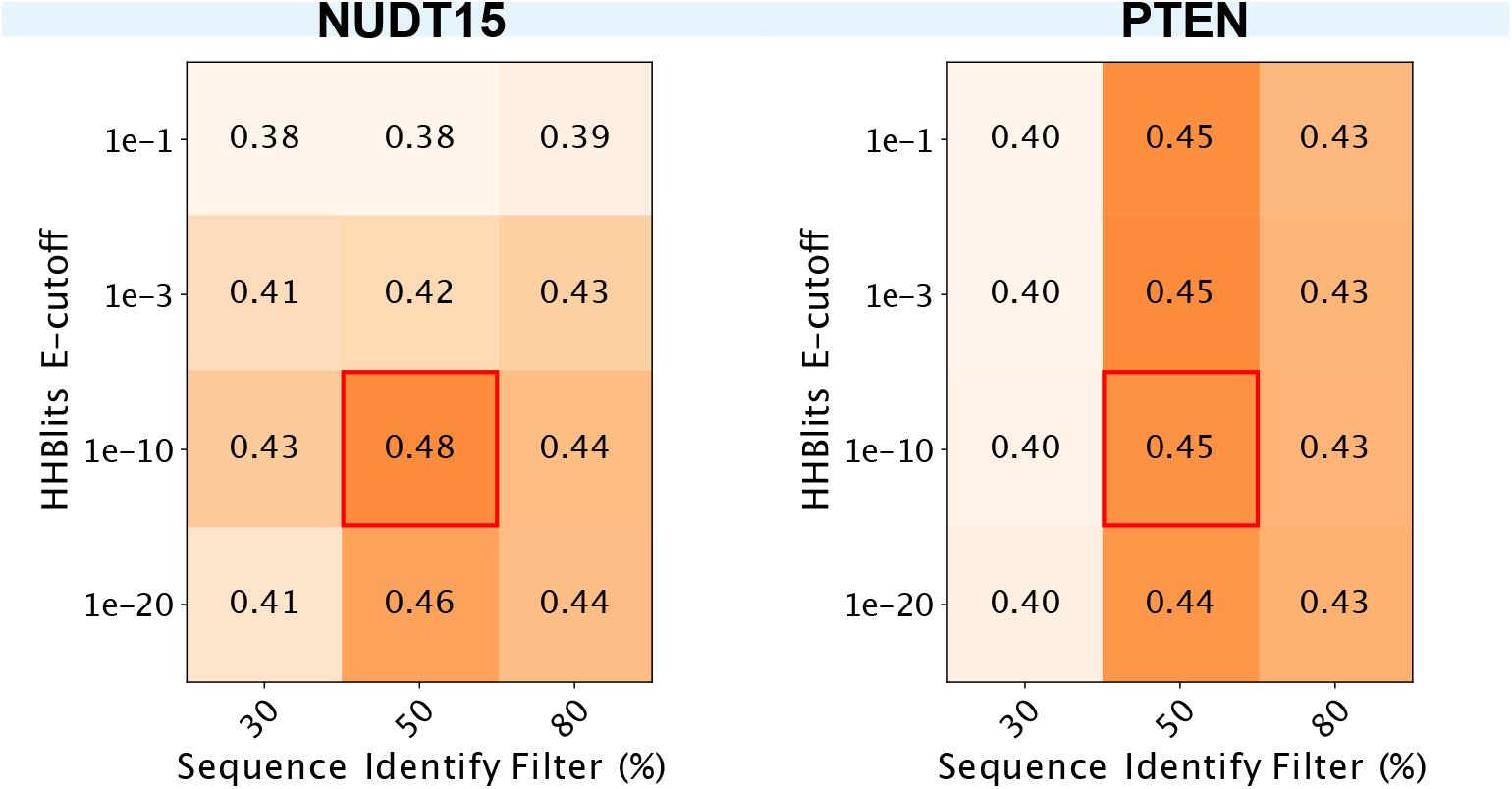
Effect of parameters used in the generation and analysis of multiple sequence alignments. We used different values of the E-value cutoff used in HHblits and the sequence identify cutoff used to filter the multiple sequence alignment in the lbsDCA calculation. For each combination of the parameters we calculated the correlation between the ΔΔ*E* scores and the scores from the activity-based MAVEs. The boxes highlighted in red are the default values used in our analysis in the main text that we chose prior to the analysis presented here.

